# Analysis of dendritic input currents during place field dynamics

**DOI:** 10.1101/2025.07.28.667243

**Authors:** Bence Fogel, Balázs B Ujfalussy

## Abstract

Neuronal activity is driven by the complex interplay between various membrane currents, often located in distinct domains of the spatially extended dendritic tree. How the effect of these currents propagates to the soma and contributes to neuronal output under in vivo conditions is not fully understood. Here, we develop a new method to measure and visualise the contributions of individual membrane currents to the somatic response in spatially extended biophysical model neurons. Our approach relies on the iterative decomposition of the axial current flowing between neighbouring compartments in proportion to the underlying membrane currents measured in the model. We apply this method to visualise the inputs driving hippocampal place cell activity. Our method provides a compact and intuitive description of the various dendritic events underlying subthreshold activity, spiking, or burst firing. By contrasting the dendritic input currents preceding spiking and bursting, we demonstrate that both could occur at highly variable input levels to proximal dendrites (basal and oblique), and that strong distal inputs facilitates, rather than controls, the generation of complex spike bursts. Our method opens a novel window onto single-neuron computations that will help to design better models and to interpret the results of in vivo imaging experiments.

## Introduction

Characterizing the relationship between the dynamics of individual ion channels and the neuronal activity *in vivo* is crucial for mechanistic understanding of biological computation, to develop more realistic circuit models of brain activity, and to identify potential therapeutic targets.

Ion channels have been extensively studied both in isolation (Hille, 2001) and also in their interactions with other channels to explain the emergence of diverse neuronal activity patterns observed (Hodgkin and Huxley, 1952; Koch, 1999). A dominant view emerged that neurons form non-linear dynamical systems, with state variables corresponding to membrane potential (Vm), open probability of ion channel gates, and second messenger concentrations (Izhikevich, 2007). Even if the membrane currents are known, studying the impact of particular ion channels on the neuronal response in such a dynamical system under *in vivo* conditions is hindered by two major obstacles: First, synaptic inputs to neurons arrive in complex temporal patterns often engaging a combination of ion channels that interact with each other in potentially highly nonlinear, state-dependent manner. Measuring channel-contributions under *in vitro* conditions (Losonczy and Magee, 2006) or by pharmacological manipulations (Palmer et al., 2014) may not engage the ion channels in the same way as during the high-conductance state *in vivo*, (Destexhe et al., 2003; Ujfalussy et al., 2018). In response to these challenges, the currentscape technique has recently been developed to provide an intuitive and simple way to visualize the high-dimensional current dynamics during naturalistic neural activity in one-compartmental biophysical neuron models (Alonso and Marder, 2019).

Second, synaptic inputs target the spatially extended dendritic tree of the neurons (Fig. 1B) where electrical and biochemical compartmentalization renders state variables local (Stuart and Häusser, 2001; London and Häusser, 2005; Branco and Häusser, 2010). With local state variables, the recruitment of ion channels can vary greatly between different dendritic domains (Häusser and Mel, 2003; Poirazi et al., 2003), which makes it especially difficult to understand or visualise current-interactions in neurons with a large dendritic tree under *in vivo* conditions. There are three simple ways to adapt the currentscape technique to neurons with realistic morphology: First, one could visualize the current dynamics in each compartment separately (Guet-McCreight and Skinner, 2020; Linaro et al., 2022). This strategy could work for few compartmental models but is clearly not feasible beyond a critical size and does not provide an intuitive description of the current propagation in neurons. Second, it is possible to sum up all membrane currents throughout the dendritic tree (Fig. 1A-D). However, this approach can not reveal the contribution of currents to the output of the cell, as even large-amplitude distal dendritic currents often fail to propagate to the soma (white arrowhead in Fig. 1D). Third, one could focus on a single compartment and study the input currents locally, for example at the somatic action potential generation site (Linaro et al., 2022). However, the dominant inputs are often currents flowing axially as a consequence of the activation of synaptic or intrinsic currents in other compartments (Fig. 1D). Here, we develop a computational technique, the *extended currentscape*, which is able to identify how membrane currents measured throughout the dendritic tree contribute to the axial current flowing into any particular compartment.

**Fig. 1:**
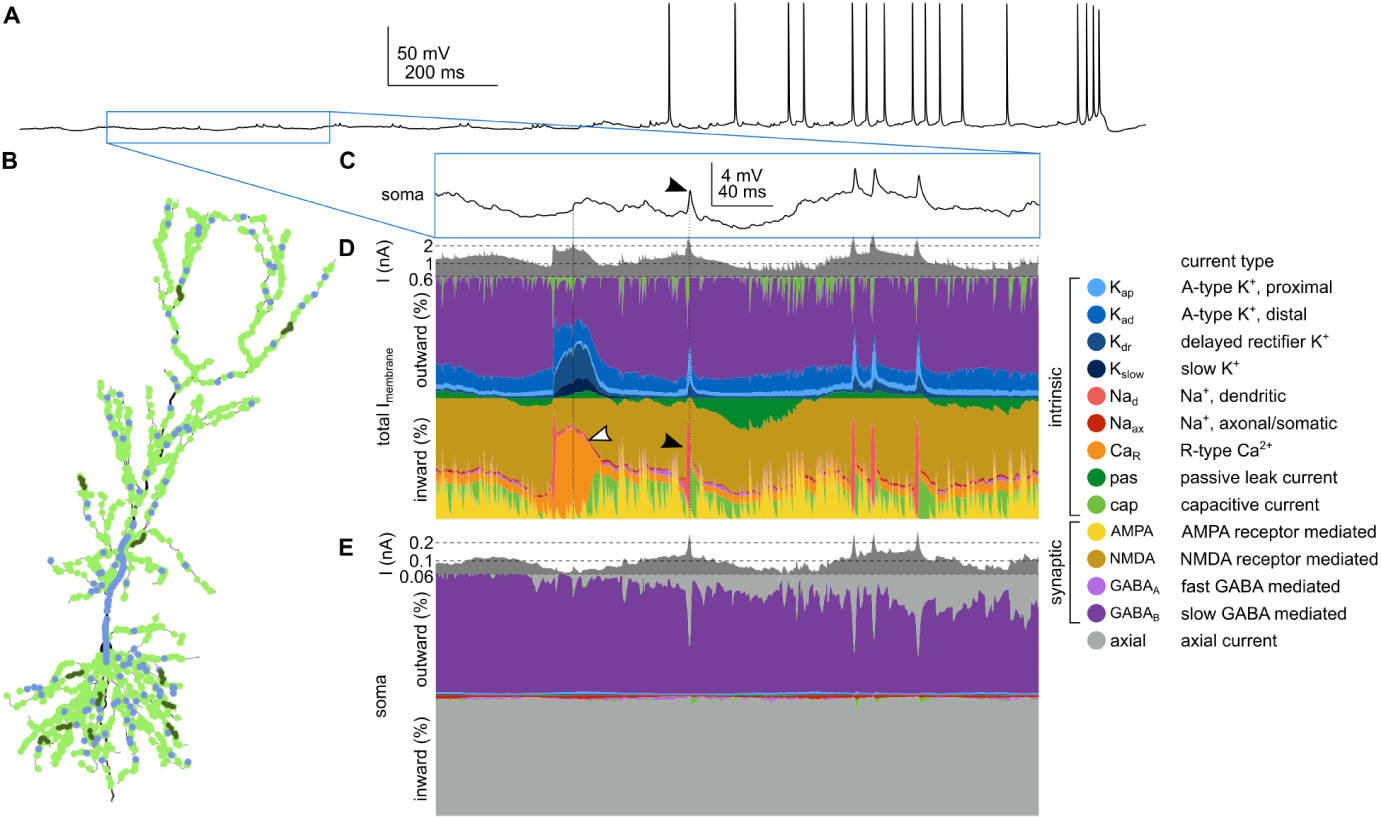
Challenges in identifying the biophysical factors underlying neural responses. **A**) Somatic Vm response of a biophysical model CA1 pyramidal neuron to distributed naturalistic synaptic inputs. Blue box highlights the portion analyzed in panel C. **B**) Morphology of the simulated neuron with the location of the synapses (green: excitatory; blue: inhibitory synapses; dark green indicates the location of 12 functional synaptic clusters. See Methods and Fig. 4 for more details.) **C**) Magnified part of the somatic Vm response. Filled arrowhead highlights a spikelet. **D**) Visualising the input currents in the model using the currentscape technique (Alonso and Marder, 2019). Top: the magnitude of the total inward current on a logarithmic scale. Since we included the capacitive current to the sum (Eq. (1)), the magnitude of the inward and the outward currents is identical (Kirchhoff’s law). Here membrane currents across the entire dendritic tree were summed. Bottom: Percentage of the different ion channels, including intrinsic and synaptic channels, contributing to outward (inhibitory, top) and inward (excitatory, bottom) currents. Color legend is shown on the right and applies to all subsequent figures. White arrowhead indicates a large Ca^2+^-current that does not appear in the somatic Vm response. Filled arrow highlights dendritic Na^+^-channel activation corresponding to spikelet in C. **E**) Currentscape applied to the somatic currents including currents flowing axially from different dendritic branches (grey).

To demonstrate the ability of the extended currentscape method to provide an intuitive visualisation of input integration in neuron models with complex morphology under *in vivo*-like conditions, we apply it to study the generation of complex spike bursts (CSB) in hippocampal CA1 pyramidal neurons (PNs). In PNs CSBs are associated with Ca^2+^-spikes in the apical dendrites (Larkum et al., 1999; Grienberger et al., 2014; Magó et al., 2021) and are believed to play a critical role in learning (Payeur et al., 2019) and memory formation (Bittner et al., 2015, 2017; Grienberger and Magee, 2022; Wen et al., 2024). However, the precise role of the different input pathways in triggering Ca^2+^-spikes in these cells remained unclear: On the one hand, Ca^2+^-spikes can be readily initiated by distal dendritic, but not by somatic current injection *in vitro* (Golding et al., 1999; Takahashi and Magee, 2009), the timing of the Ca^2+^-spikes coincides with inputs to the most distal branches during theta oscillation (Bittner et al., 2015) and blocking entorhinal inputs decreases the probability of CSB firing (Bittner et al., 2015) all emphasizing the critical role of distal inputs in controlling CSBs. On the other hand, CSBs can be triggered by somatic current injection in most PNs at random spatial locations *in vivo* (Bittner et al., 2015, 2017) or *in vitro*, after blocking dendritic potassium channels (Bittner et al., 2017), distal inputs alone are not as effective as combined with proximal inputs in triggering CSBs (Takahashi and Magee, 2009) and distal tuft dendrites are variably recruited during putative CSB events *in vivo* (O’Hare et al., 2025), raising the possibility that proximal inputs also play an important role in the initiation of CSBs.

**Table 1:**
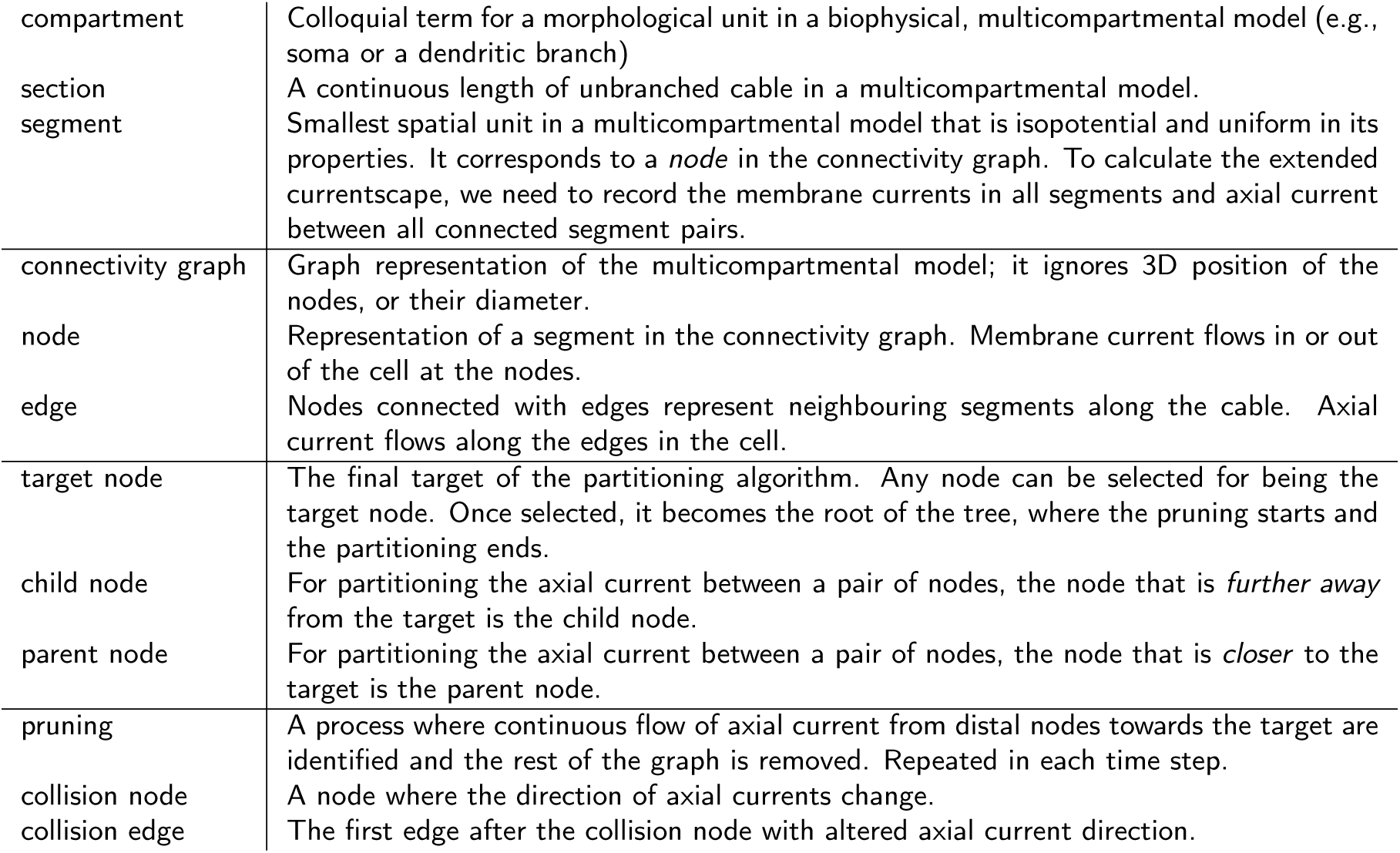
Glossary.

|  |  |
| --- | --- |
| compartment | Colloquial term for a morphological unit in a biophysical, multicompartmental model (e.g., soma or a dendritic branch) |
| section | A continuous length of unbranched cable in a multicompartmental model. |
| segment | Smallest spatial unit in a multicompartmental model that is isopotential and uniform in its properties. It corresponds to a *node* in the connectivity graph. To calculate the extended currentscape, we need to record the membrane currents in all segments and axial current between all connected segment pairs. |
| connectivity graph | Graph representation of the multicompartmental model; it ignores 3D position of the nodes, or their diameter. |
| node | Representation of a segment in the connectivity graph. Membrane current flows in or out of the cell at the nodes. |
| edge | Nodes connected with edges represent neighbouring segments along the cable. Axial current flows along the edges in the cell. |
| target node | The final target of the partitioning algorithm. Any node can be selected for being the target node. Once selected, it becomes the root of the tree, where the pruning starts and the partitioning ends. |
| child node | For partitioning the axial current between a pair of nodes, the node that is *further away* from the target is the child node. |
| parent node | For partitioning the axial current between a pair of nodes, the node that is *closer* to the target is the parent node. |
| pruning | A process where continuous flow of axial current from distal nodes towards the target are identified and the rest of the graph is removed. Repeated in each time step. |
| collision node | A node where the direction of axial currents change. |
| collision edge | The first edge after the collision node with altered axial current direction. |

In this paper, we first describe the *extended currentscape* technique that generalizes the standard currentscape visualization method to neurons with spatially extended dendritic trees. We show that the extended currentscape accurately and intuitively captures the origin and type of dendritic events in models with simplified or realistic morphology under spatially localised synaptic stimulation conditions. Next, we analyse the membrane currents and the Vm dynamics of a biophysical model CA1 PN showing place cell-like activity. We show that the membrane potential throughout the entire dendritic arbor has low-dimensional dynamics dominated by global activity even when localised dendritic events are abundant in distal dendrites. Next, we analyze CSBs in the model using the extended currentscape method and show that they can be started with highly variable initiation dynamics. By contrasting CSBs and isolated single spikes, we reveal that although distal inputs facilitate the generation of CSBs, there is no need for exceptionally strong tuft inputs nor Ca^2+^ hotspots for their initiation.

## Results

### The extended currentscape method

We start by describing the extended currentscape visualization method which aims to capture how membrane currents in a distant *reference* (e.g., dendritic) compartment influence the activity of a *target* (e.g., soma) compartment. Our method relies on the equivalent circuit model of neurons (Koch and Segev, 2000) and requires measuring the membrane and axial currents throughout the dendritic tree of a neuron (in every node of the circuit) to partition the axial currents flowing into the target compartment by the underlying membrane currents. Since this kind of data is not available in real neurons, we used multicompartmental biophysical models (Jarsky et al., 2005; Ujfalussy and Makara, 2020) to develop and test our method. With the advent of novel single-neuron voltage imaging techniques (Brooks et al., 2024; Liao et al., 2024; Park et al., 2025; Lee et al., 2026), our method could potentially also be adapted to data from real neurons in the near future.

The intuition behind our method is that distant currents can directly influence the Vm of the target compartment only if there is a continuous flow of axial current from reference to target, i.e., if the axial current is not blocked or reversed between the reference and the target. In order to characterise the relationship between the axial currents and the membrane currents we start with Kirchhoff’s current law, stating that the sum of all currents flowing into or out of a node in an electrical circuit must be zero. Applying it to a small segment of a neuronal process, we have:

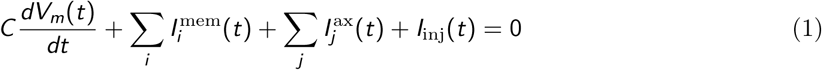

Here *C* is the membrane capacitance, 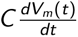 is the capacitive current, *I* ^mem^ denotes the membrane currents, including all synaptic and intrinsic currents, *I*^ax^ denotes all axial currents flowing from the neighbouring segments and *I*_inj_ is the current injected into the cell through a stimulation electrode. Eq. (1) states that if the axial current flows from the target towards a child node (as at t_1_ in Fig. 2A), then the sum of the outward currents must be larger than the sum of the remaining inward currents in the child compartment. In this case, we postulate that this excess outward current is responsible for the axial current flowing towards the child compartment, and we partition the axial current in the proportion of the outward currents in the child compartment (Fig. 2B, left). In contrast, when the direction of the axial current between the child and the target reverses (Fig. 2A, t_2_), we partition it proportionally to the inward currents in the child compartment (Fig. 2B, right). Note that the sign of membrane currents can also reverse when the membrane potential crosses their reversal potential. In our simulations this often happened with the leak current, which had its reversal potential (*E*_leak_ = −66 mV) between that of excitatory (*E*_NMDA_ = 0 mV) and inhibitory inputs (*E*_GABA_ = −80 mV). This way, the same current type can contribute to inward or outward axial currents at different time points or in different compartments.

**Fig. 2:**
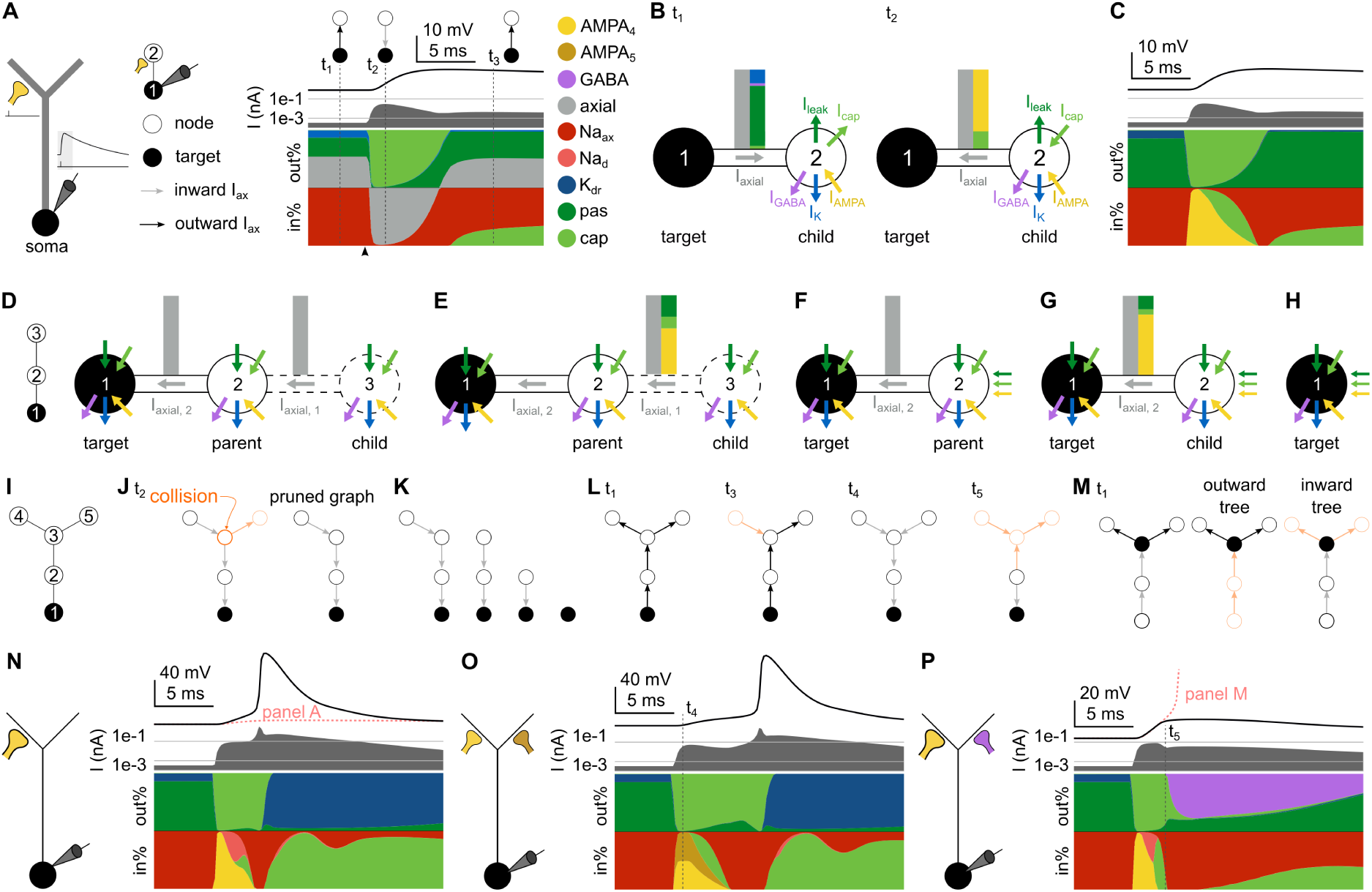
Partitioning the axial current. **A** Currentscape analysis of a simple model responding to excitatory synaptic input. Left: Morphology of the model with a soma, an apical trunk and two dendritic branches. Insets show the stimulus and the somatic response, with the period analysed in later panels highlighted. Middle: simplified graph representation of the model with two nodes. Here we used this 2-node graph only for illustration purposes. See panels D and I for larger graphs describing the same biophysical model. Right: currentscape of the somatic compartment of the model. Vertical dotted lines indicate the time points analysed in panels B, J and L, with the corresponding axial current directions shown as insets; arrowhead show the timing of the input. pas: passive leak current. cap: capacitive current. **B**) The axial current of the *target* compartment (grey arrow) is partitioned by the membrane currents in the *child* compartment (coloured arrows). When the axial current flows away from the target, it is partitioned by the outward currents in the child (t_1_, left). When the direction of the axial current reverses, it is partitioned by the inward currents in the child (t_2_, right). **C**) Extended currentscape of the somatic node shown in panel A. **D-H**) Partitioning recursion. Partitioning begins at the most distal child compartment (3) and moves through its parent towards the target (e.g., 1–soma; D). In the first step, *I*_axial,1_ is partitioned proportionally to the inward currents in node 3 (E). Next, the partitioned axial currents are added to the membrane currents in the parent (node 2; F). Steps D-F are repeated for the next pair of nodes (F-G) until the target is reached. **I-L**) Iterative partitioning in a graph. **I**) Graph representation of a multicompartmental model. The compartments (nodes) are denoted by circles and the axial current flow is indicated by arrows (edges). **J**) The graph, representing the flow of axial currents at t_2_ in panel A is pruned at the *collision* edges, where the direction of the axial current reverses. **K**) Partitioning algorithm starts from a leaf node and progresses towards the target. **L**) The structure of the pruned graph is time dependent: each graph shows a pattern of axial current flow at different time points from panels A, O or P. Orange colours highlight the part of the graph behind the colliding edge that can not directly contribute to axial currents towards the target. **M**) Partitioning with node 3 as the target: inward and outward currents are considered separately for both pruning and partitioning. **N-P**) Currentscape analysis of the responses of the simple model to strong synaptic input evoking a dendritic Na^+^-spike triggering a somatic AP (N), coincident excitatory inputs to different dendritic branches (O) and inhibitory input blocking somatic AP after a dendritic Na^+^-spike (P).

Fig. 2A-C illustrates the application of the extended currentscape to a simple biophysical model in which synaptic input to the dendrite evokes an EPSP that propagates to the soma. As we want to analyse the currents driving the soma, we choose the soma to be that target compartment throughout the analysis (the user can pick any compartment to be the target but it remains the target throughout the analysed time period). In this model, small baseline Na^+^-current makes the soma more depolarized than the dendrite in the equilibrium, and the axial current flows outward from the target compartment (Fig. 2A-B, t_1_). After the input is activated, the axial current reverses and depolarizes the soma giving rise to the measured EPSP (Fig. 2A-B, t_2_). During the repolarization phase (Fig. 2A, t_3_) the axial current returns to its original direction flowing from the soma towards the dendrites. The extended currentscape plot reveals that the somatic depolarization was mainly caused by synaptic AMPA receptor mediated currents (yellow in Fig. 2C), while the outward currents left the cell via leak currents (dark green in Fig. 2C).

In a multicompartmental model, the partitioning of the axial currents can be applied iteratively starting from the most distal compartment and proceeding towards the target once all axial currents arriving at the parent from the more distal nodes had been partitioned (Fig. 2D-E). In the next step, the partitioned axial currents are added to the membrane currents in the parent node (Fig. 2F) before moving one step closer to the target (Fig. 2G-H).

In general, the topology of the model is represented by an acyclic graph, where each node is a segment, the directed edges represent the flow of the axial current and the root node of the graph is the target (Fig. 2I-L). Partitioning starts with following the flow of axial currents from the target towards the leaf nodes of the tree. The nodes where the direction of the axial current is reversed are collision points (Fig. 2J). Since the propagation of the axial current is blocked at the collision points, the membrane currents of the nodes distal to the colliding edge do not have a direct influence on the activity at the target. Therefore, to make the algorithm computationally more efficient, in every time step we prune the graph at collision edges and apply the partitioning recursion only to the pruned graph (Fig. 2J-K). Since the direction of the axial currents often changes during the simulations, the shape and size of the remaining graph can vary between different time points (Fig. 2L). The target node may connect multiple subtrees, and the axial currents of the subtrees are pruned and partitioned separately (Fig. 2M).

After partitioning the axial currents of the target compartment we can use the standard currentscape plot (Alonso and Marder, 2019) to visualize the contribution of distal membrane currents to the membrane potential dynamics of the target. The entire process must be repeated when a new target compartment (e.g. a dendritic branch) is selected (Fig. 2M). Throughout the paper, we will use two different variants of the partitioning algorithm: we will either partition axial currents by the *type* of membrane currents (e.g., current flowing through Ca^2+^ or Na^+^ channels) or by the dendritic *region* of the reference compartments (i.e., basal, oblique or tuft dendrites). However, alternative variants of the partitioning algorithm can also be devised (see Methods).

To illustrate how the extended currentscape analysis can provide a compact summary of the distal currents driving the somatic response in simple, intuitive situations, we show three additional examples (Fig. 2M-O). First, we increased the distal synaptic conductance, which now evoked a local dendritic Na^+^-spike (Fig. 2M), and the somatic EPSP became large enough to trigger an action potential (AP). Similarly, if two weak synapses are stimulated coincidentally at different dendritic branches, their combined activation can also lead to a somatic AP (Fig. 2N). Although the axial currents arriving to the soma are similar in these cases, the extended currentscape can reveal the differences in the membrane dynamics leading to the somatic spikes. Finally, off-the-path inhibition can prevent the generation of the somatic AP triggered by the local dendritic spike (Fig. 2O). In this case, the strong, GABA-receptor mediated outward currents are visible on the currentscape plots during the repolarization phase of the EPSP.

Next, we will apply the extended currentscape technique to analyze input integration and the mechanism of burst firing in hippocampal place cells.

### Currentscape analysis of dendritic integration in the CA1 pyramidal neuron model

To study the synaptic input conditions that lead to burst firing under *in vivo*-like conditions we used a previous CA1 PN model containing Na^+^-channels, a delayed rectifier K^+^ channels and two variants of A-type K^+^-channels in basal and apical dendrites (Jarsky et al., 2005; Ujfalussy and Makara, 2020) and extended it with a minimal set of channels necessary for the generation of complex spike bursts. Experimental data indicates that R-type Ca^2+^-channels are mainly responsible for both burst firing and for dendritic plateau potentials in this cell type (Magee and Carruth, 1999; Metz et al., 2005; Takahashi and Magee, 2009). We used a novel Ca^2+^-channel model that displayed similar activation and inactivation kinetics to the Ca^2+^ currents recorded in the apical dendrites in CA1 PNs (Fig. S1A; Magee and Johnston 1995) and distributed it uniformly across all apical dendrites (trunk, obliques, and tuft) of the cell (Magee and Johnston, 1995; Poirazi et al., 2003).

Various potassium channels contribute to the repolarization after Ca^2+^-spikes, including Ca^2+^-activated potassium channels (Golding et al., 1999; King et al., 2015), but the precise biophysical characterization of these channels is currently lacking. To model their overall effect, we used a high voltage activated potassium current with relatively slow kinetics (Fig. S1B) in the apical dendrites.

Equipped with these channels, our model was able to fire a short burst of 3-4 action potentials with decreasing amplitude riding on a depolarizing wave upon dendritic current injection (Fig. S1C). These somatic bursts were accompanied by dendritic Ca^2+^-spikes, so we refer to them as complex spike bursts or CSBs. In agreement with the experimental data (Golding et al., 1999) the model had a lower threshold for initiating Ca^2+^-spikes in the apical trunk than in the soma (Fig. S1D). Under *in vivo*-like synaptic input conditions (see below and Methods), dendritic Ca^2+^-spikes could also be evoked by somatic current injection (Fig. S1E), as in Bittner et al. (2015).

To illustrate the ability of the extended currentscape method to provide a compact and intuitive summary of dendritic events in the model, we first tested it under simple, spatially and temporally restricted input conditions. We first used the model without dendritic Ca^2+^-channels and stimulated an increasing number of synapses in a single oblique branch Fig. 3A-C). We found that the somatic response became superlinear beyond a branch-specific threshold, with a sigmoid superlinearity (Fig. 3A-B), reminiscent of experimental findings with 2-photon glutamate uncaging (Losonczy and Magee, 2006). Partitioning the somatic currents by the current type (Fig. 3C, middle) indicated that before stimulation, the inward currents were mainly leak currents (dark green), with some contribution from the somatic and axonal Na^+^-currents (red). Partitioning the somatic currents by the region of their origin (Fig. 3C, bottom) showed, that current flew from the axon and basal dendrites towards the soma and further towards the apical dendrites, that were more hyperpolarized than the soma due to the larger density of K^+^ channels in the apical trunk. Upon stimulation, the inward currents show first a fast, AMPA-mediated and later a slower NMDA-mediated component (yellow and olive in Fig. 3C, middle), both originating from the stratum radiatum (levander in Fig. 3C, bottom). At n=20 inputs a Na^+^-spike that remains local to the stimulated branch is responsible for the spikelet appearing in the somatic response (pink in Fig. 3C, right). During these events, the inward current arrived transiently from the apical dendrites (see shades of levander in Fig. 3C, bottom), and left the soma towards the basal dendrites (pink in Fig. 3C). At the end of the events the NMDA contribution disappeared from the soma as the axial current reversed to its original direction, flowing from basal towards apical dendrites.

**Fig. 3:**
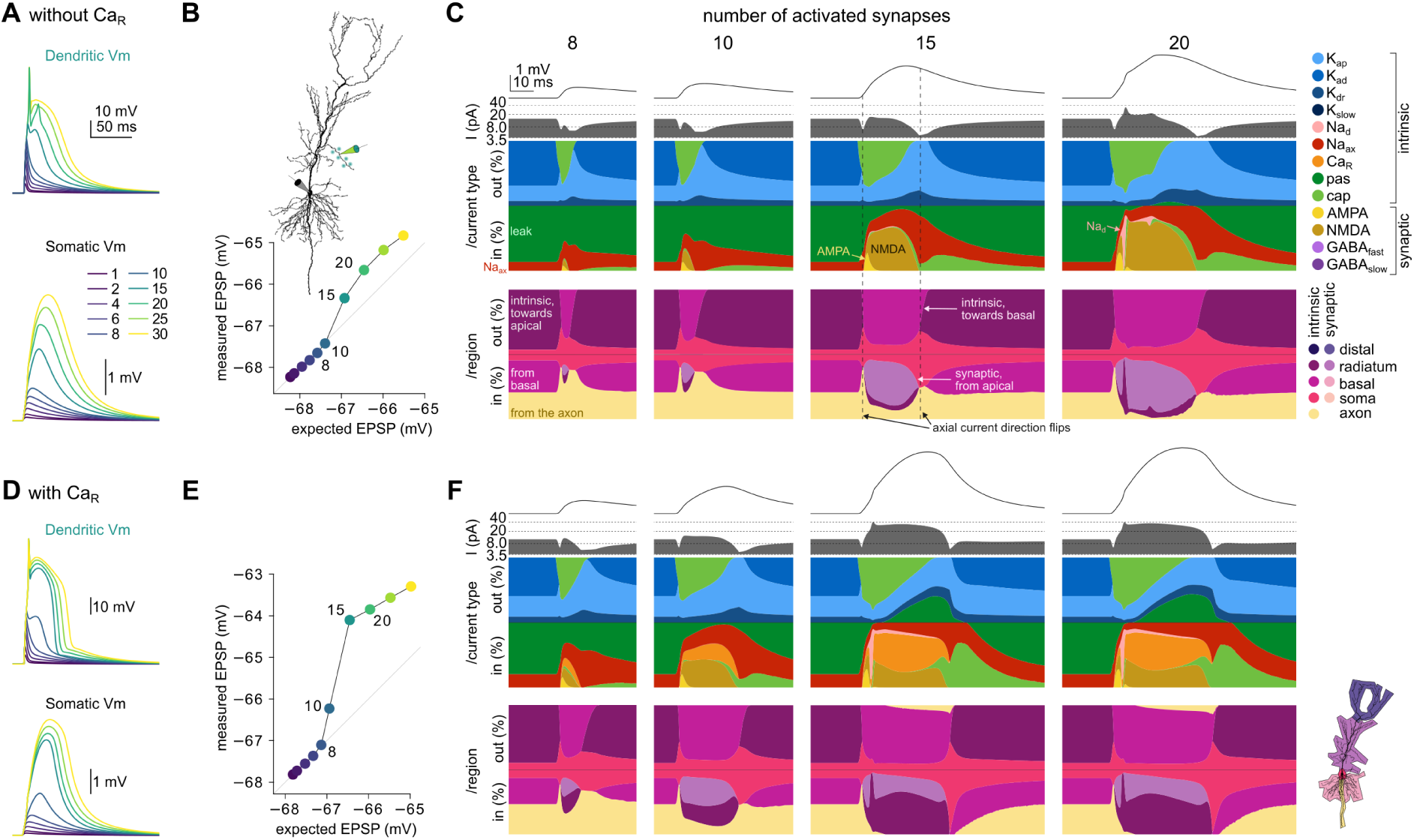
Currentscape analysis of dendritic integration in the CA1 PN model. **A**) Dendritic (top) and somatic (bottom) Vm in response to stimulating an increasing number of synapses (N=1-30) on an oblique dendrite (inset in B) with 0.3 ms delay in the model without Ca^2+^channels. Note the fast dendritic Na^+^-spike appearing at n=20. **B**) Expected versus measured somatic response amplitude of the stimulations shown in A. Inset shows the branch used for stimulation and dendritic recordings. **C**) Extended currentscape analysis of the somatic responses to an increasing number of stimulations (n=8, 10, 15 and 20 shown). Top line: somatic membrane potential (Vm) response. Second line: total outward membrane current on log-scale. Third line: percentage of somatic outward and inward currents partitioned by the current type. Fourth row: somatic currents partitioned by the current origin. **D-F**) Same as A-D for the model equipped with Ca^2+^-channels in the apical dendrites. Note the step-like response in the dendritic Vm (D, bottom) and the large Ca^2+^-currents for n=15 and 20 stimuli (F, right).

Next we repeated the same stimulation protocol after including the Ca^2+^-channels to the apical dendrites of the neuron. The addition of the Ca^2+^-channels rendered the shape of the superlinearity more step-like, as in single-photon uncaging or synaptic stimulation experiments (Fig. 3D-E; Wei et al. 2001; Ariav et al. 2003; Cai et al. 2004). Under these conditions the combined Ca^2+^- and NMDA-spikes did not lead to somatic action potential firing but remained localised to the stimulated branch. The currentscape analysis revealed that activation of Ca^2+^-channels first boosted the NMDA currents activated at near-threshold responses (at n=10 inputs, orange in Fig. 3F), and were further amplified by local Na^+^-spikes (at n=15 or 20 inputs; Fig. 3F). During these events, NMDA-mediated and Ca^2+^-currents contributed similarly to the somatic depolarization (Fig. 3).

Taken together, these simulations confirmed that extended currentscapes provide a rich form of visualization of the sequence of dendritic events leading to somatic responses even in cells with complex morphology under relatively simple input conditions. Next, we used this model to investigate the synaptic input patterns leading to CSB firing during *in vivo*-like inputs.

### CSBs in the CA1 pyramidal neuron model

To create place cell activity in response to *in vivo*-like input condition we simulated the activity of 2000 excitatory and 200 inhibitory presynaptic neurons during the traversal of a 2 m long linear track as described previously (Fig. 4A; Ujfalussy and Makara 2020; Kim et al. 2023). Briefly, each excitatory neuron had a ∼ 20 cm long place field and neurons showed theta modulation and displayed phase precession (Skaggs et al., 1996). Place fields were distributed uniformly across the linear track. The majority of the 2000 excitatory synapses were placed randomly on the dendritic tree (avoiding the soma and the proximal apical trunk; light green in Fig. 1B), while the remaining 240 inputs active in the middle of the track were organized into 12 functional synaptic clusters (Adoff et al., 2021) where presynaptic neurons with similar place fields (dark green circles in Fig. 4A) innervated neighbouring dendritic locations (Fig. 1B; Adoff et al. 2021) and had larger synaptic conductance (see Methods; Bittner et al. 2015; Heredi et al. 2025). These synaptic clusters provided strong drive to the cell in the middle of the track by activating a few dendritic branches (Ujfalussy and Makara, 2020).

**Fig. 4:**
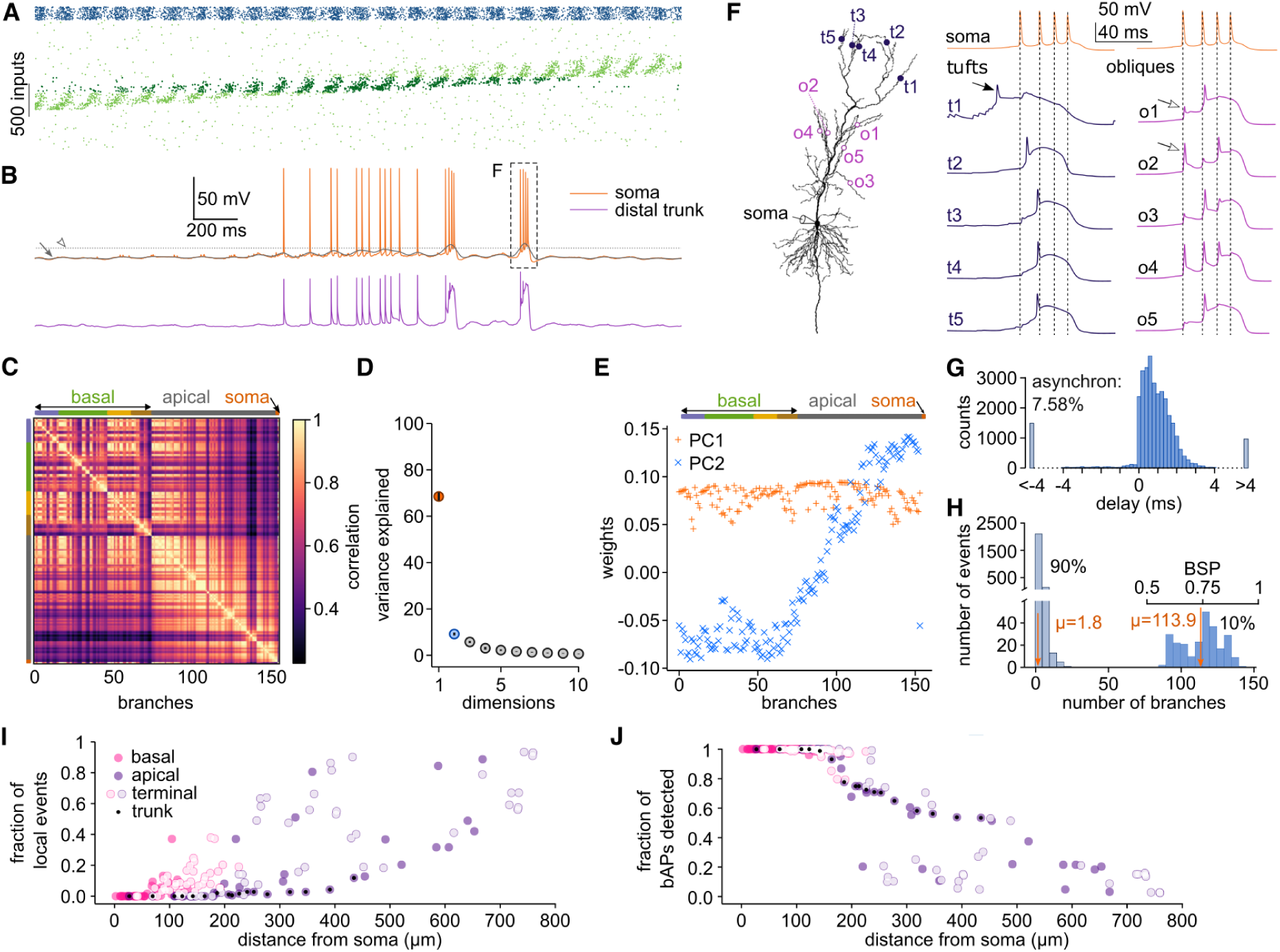
Model response to complex synaptic inputs. **A**) Activity of the 2000 excitatory (green; ordered by the place field location) and 200 inhibitory synapses (blue). Note the theta oscillation and the theta sequences in the excitatory inputs. Synapse locations are shown in Fig. 1B. Dark green indicates the 240 excitatory synapses with stronger weights and organized to functional clusters (see Methods). **B**) Somatic (top) and dendritic (from distal apical trunk, bottom) Vm response of the model to the synaptic input pattern shown in A. Grey line (arrow) shows the filtered somatic Vm response used to detect complex spike bursts (CSB). CSBs coincided with large depolarizing events in the dendrite. Dotted line (empty arrowhead): CSB detection threshold. The second CSB event (box) is analyzed further in panel F. **C**) Average (n=16 laps) cross-correlation of the Vm of the 153 dendritic branches in the model. Branches are ordered by morphology of the cell, not by correlation strength. Soma is shown as branch 153. **D**) The correlation matrix is low rank: the first two components (orange and blue) explain ∼80% of the variance of the Vm. Error bars show SD across 16 simulations. **E**) The weights of the first two principal components (PC1 and PC2) in an example simulation: the first component describes a uniform activity across the entire cell, while the second captures activity differences between the perisomatic region and distal apical dendrites. **F**) Left: location of the dendritic Vm recordings (t: tuft, o: oblique, s: soma). Right: Dendritic Vm during somatic burst firing. Filled arrow indicates a local dendritic Na^+^ spike, open arrows highlights backpropagating APs. **G**) Histogram of the time difference between dendritic Vm peaks (reaching -30 mV, with a prominence of 30 mV) and the closest Vm peak at the soma, calculated from 16 simulations and the 153 dendritic branches. **H**) Histogram of the spatial extent of each dendritic event. The 90% of the events were local (light blue), involving *<* 20 branches (*µ* = 1.8). The remaining 10% (dark) were associated with somatic APs. Branch spike prevalence (BSP) of these events is shown on the top. **I-J**) Fraction of local events (I) and fraction of bAPs detected as a dendritic event (J) as a function of the distance of the branch from the soma. Light color: terminal branches; black dot: apical trunk.

Inhibitory neurons showed theta modulation (Csicsvári et al., 1999) but were spatially untuned (Wil-son and McNaughton, 1993; Dupret et al., 2013) consistent with their indirect role in shaping spatial selectivity in place cells (Grienberger et al., 2017). Half of the inhibitory cells targeted the soma and the proximal apical trunk, the other half was randomly distributed throughout the dendritic tree (blue circles in Fig. 1B; Megias et al. 2001).

First, we analyzed the somatic and dendritic Vm response of the model. The neuron responded to synaptic stimulation with a train of action potentials in the middle of the simulated track consisting of a variable number of single spikes and CSBs (Fig. 4B). During CSBs the distal apical dendrites showed a Vm response similar to the Ca^2+^-spikes observed *in vitro* (Fig. 4B, bottom trace). In a set of 16 simulations with variable cluster placement and synaptic input patterns, the Vm of the entire neuron was highly correlated (Fig. 4C), with a single factor explaining ∼70% of the total variance across the dendritic branches (Fig. 4D, Lee et al. 2026). The weights associated with this first factor uniformly covered the entire dendritic tree (Fig. 4E, see also Fig. S2) indicating the dominance of the global modulation of the dendritic Vm mainly due to back-propagating action potentials (bAPs, Spruston et al. 1995). In contrast, the weights associated with the second principal component had an opposite sign perisomatically versus distally, demonstrating a greater level of independence from the soma in distal tuft branches (Fig. 4E).

When we inspected the dendritic Vm during CSBs we found that in our model CSBs were not preceded by large, global Ca^2+^-spikes in the tuft. Instead, dendritic Ca^2+^-spikes typically started asynchronously (O’Hare et al., 2025), often after a back-propagating action potential or a local Na^+^-spike (Fig. 4F, see more examples in Fig. S3; Park et al. 2025; Lee et al. 2026). Although in distal dendrites we could observe local Na^+^ or Ca^2+^-spikes not coupled to somatic activity (see below in Fig. 5), local dendritic Vm peaks closely followed somatic APs in the majority of cases (>90%; Fig. 4G). When we analysed the spatial extent of the dendritic events, we found two distinct clusters Fig. 4H): most (90%) of the events were local, restricted to a small subtree (*<* 20, *µ* = 1.8 branches). These local events occurred in higher-order (mostly terminal) basal or apical branches (Fig. 4I). The second group was associated to bAPs reaching the majority of the dendritic branches, with propagation failures occurring mainly in higher-order apical dendrites beyond 200 *µ*m from the soma (Fig. 4J; Lee et al. 2026). This way, although only 10% of the dendritic Vm events were associated with bAPs, they reached ∼60-times more branches than local events and they dominated the PCA analysis even in the presence of local regenerative dendritic events driven by strong, functionally clustered synaptic inputs.

**Fig. 5:**
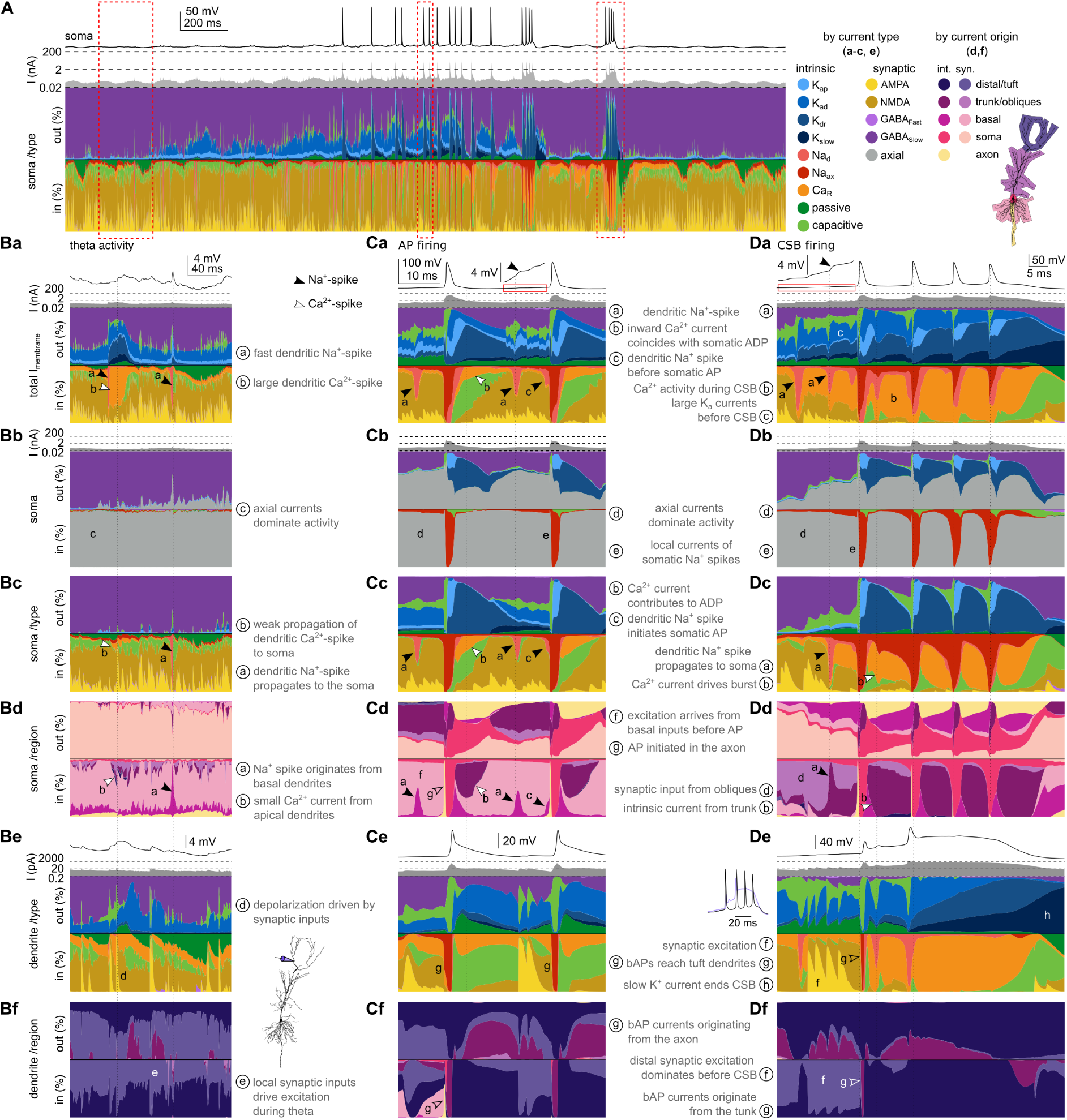
Current dynamics in a place cell. **A**) From top to bottom: Somatic Vm; inward somatic currents (log scale) and percentage of different outward and inward currents. Colour legend is shown on the right. Red boxes highlight temporal intervals analyzed in panels B-D. Filled arrowheads in panels B-D indicate local dendritic Na^+^-spikes, open arrowheads point to bAPs and white arrowheads highlight Ca^2+^-spikes. **B**a-f) Current dynamics outside of the place field. **B**a) Total membrane current of the cell during theta oscillation. **B**b) Somatic currents, including currents flowing from dendrites (axial currents, grey). **B**c) Somatic currents partitioned by current type. White arrowhead highlights that dendritic Ca^2+^-spike has very little contribution. **B**d) Somatic currents partitioned by current source region. Colour legend is shown on the right of panel A. Arrowheads highlights that Ca^2+^-spike originated in the apical region whereas the Na^+^-spike came from a basal dendrite. **B**e) Membrane potential (top), sum of inward currents (middle) and input current types in a tuft branch. The recording location is shown as an inset next to panel Be. **B**f Tuft currents partitioned by current region. **C**a-f) Same as panel B during action potential (AP) firing. Dendritic Na^+^-spikes originating in the basal dendrites (Cd) appear as spikelets in the soma (filled arrowhead; inset in Ca). The cell is mostly driven by NMDA inputs (Cc) targeting basal dendrites (Cd) and the tuft is largely decoupled from the rest of the neuron, though APs back-propagate (open arrowhead in Cf). **Da-Df**) Same as panel B during CSB firing. CSB is preceded by multiple dendritic Na^+^-spikes, some propagating to the soma (black arrowhead in Dc) from stratum radiatum dendrites (Dd). This facilitates a Ca^2+^-spike from the oblique-trunk region to propagate to the soma (Da-Dd) leading to the first somatic AP. The back-propagating AP (open arrow in Df) triggers Ca^2+^-spike in tuft and oblique branches that efficiently propagates to the soma and triggers CSB (white arrowhead in Dc and Dd). Further details are in the legend.

### Currentscape analysis of place field dynamics

To better understand how individual dendritic events influenced the somatic response of the cell, we calculated the extended currentscape of the simulated neuron in the time window around its activity in the place field (Fig. 5), and analysed the membrane currents at subthreshold activity (Fig. 5B), AP firing (Fig. 5C) and during complex spike bursts (Fig. 5D). Partitioning the somatic axial currents by current type revealed that before entering the place field, the cell was driven mainly by synaptic currents (Fig. 5Ba-c). Voltage dependent Na^+^ or Ca^2+^ channels only activated occasionally and briefly, and their effect often remained localized, not being able to efficiently propagate to the soma (Fig. 5Ba-c). When the effect of dendritic Na^+^-spikes reached the soma, they either appeared as small spikelets (Fig. 5Ba) or as fast rise in the somatic Vm (Fig. 5Ca). Partitioning the somatic axial currents by their origin indicated that, in this simulation, the soma was driven mainly by the basal dendrites (Fig. 5Bd). Performing similar partitioning of the axial currents in the tuft by their type (Fig. 5Be) and by their origin (Fig. 5Bf) revealed that the tuft region was driven by local synaptic currents and was largely decoupled from the rest of the cell.

Within the place field, somatic action potentials were evoked directly by synaptic inputs (first AP in Fig. 5C) or triggered by Na^+^-spikes propagating from nearby basal dendritic branches (second AP in Fig. 5C). At the beginning of the action potential, a brief current originating from the axon can be observed (Fig. 5Cd), indicating that APs are initiated in the axon. In our model, active backpropagation of the APs is limited to the apical trunk, since the Na^+^-channels in the higher order dendritic branches have a higher voltage threshold (Ujfalussy and Makara, 2020). Indeed, back-propagating APs do not recruit local, dendritic-type Na^+^-channels in the tuft (Fig. 5Ce). In contrast, Ca^2+^-currents are recruited in the apical dendrites and contribute to the spike after depolarization (Fig. 5Ca-f).

The model neuron also fired complex spike bursts (CSBs) within its place field (Fig. 5D). When we magnified the current dynamics around the second CSB, we found that it was preceded by strong Ca^2+^-currents and dendritic Na^+^-spikes that eventually reached the soma and elicited a somatic AP (Fig. 5Da-d). In this case, the cell was driven by a mixture of synaptic and intrinsic inputs from the apical dendrites already before the CSB (Fig. 5Dd). The first AP propagated back to the dendrites and amplified the dendritic Ca^2+^- and Na^+^-currents (Fig. 5Db-d). These intrinsic currents, originating from the apical dendrite, became the dominant inward drive in the soma and drove the cell to fire further APs (Fig. 5Db-d) until the slow potassium current in the dendrites terminated the burst (Fig. 5De-f). Taken together, the extended currentscape plots provide compact and intuitive visualization of the complex current dynamics underlying neuronal activity during spatially and temporally structured synaptic inputs in single trials. Next, we set out to systematically study the conditions leading to CSB firing in our model.

### Input conditions for complex spike burst generation

To identify the input conditions leading to complex spike burst firing in our model, we focused on 41 CSB events from 16 simulations with different cluster location and input patterns, and contrasted them with 58 isolated action potentials (iAP; no other APs within 30 ms). First, we focussed on the membrane potential dynamics in long (>60 *µ*m) terminal branches in the basal, oblique or tuft region. We found that during CSBs the Vm was highly variable across events, but was typically substantially depolarized for ∼40 ms in both tuft and oblique dendrites (Fig. 6A), indicating the presence of dendritic Ca^2+^-spikes in these branches. The large depolarizations associated with local Ca^2+^-spikes were absent in basal dendrites or during iAPs (Fig. 6A-B).

**Fig. 6:**
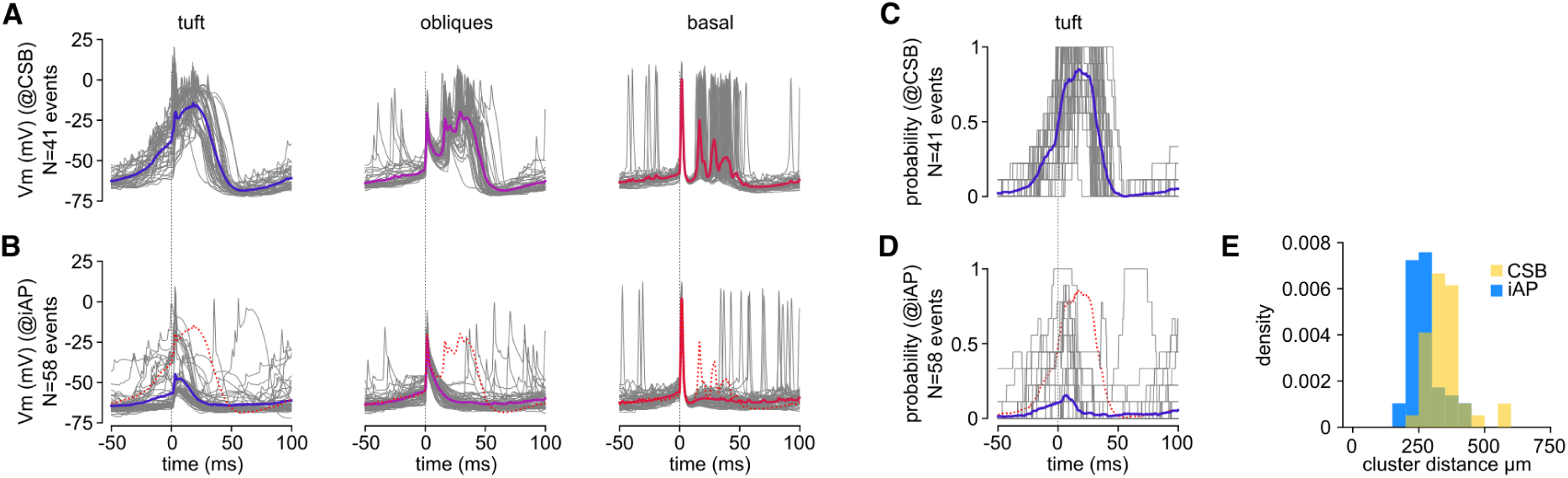
Dendritic Vm dynamics during CSBs and isolated spikes. **A**) Average dendritic Vm at somatic CSB firing. Grey lines show average across several branches in a given dendritic domain (tuft, obliques and basal dendrites) during individual CSB events; coloured lines show average across 41 CSB events. **B**) Average dendritic Vm at temporally isolated action potentials (iAPs) in different dendritic domains. Coloured lines show average across 58 events. Dotted red line shows the average response to CSB. **C-D**) Probability of Ca^2+^-spikes in tuft dendrites aligned to the start of CSBs (C) or isolated somatic spikes (D). **E**) Mean path distance of the active synaptic input clusters from the soma during CSBs and isolated spikes (t-test: *p* = 10*^−^*^5.98^).

Interestingly, the depolarization started earlier in the tuft than in the oblique branches, where it reached its maximum level only after the second AP in the burst. During CSBs most tuft branches participated in Ca^2+^-spike firing (Fig. 6C), whereas the prevalence of dendritic Ca^2+^-spikes remained low during isolated spikes (Fig. 6D). In the tuft branches, but not in obliques, both the mean Vm and the probability of Ca^2+^-spikes diverged between CSBs and iAPs already ∼10 ms before the start of the event (Fig. 6B,D). Note that the synapse density, the ion channel mechanisms and the input statistics were identical for tuft and oblique branches, suggesting that the morphology of the cell could be a primary factor underlying the increased excitability of the tuft for Ca^2+^-spikes.

To more directly test the involvement of tuft dendrites in CSB generation, we compared the location of active synaptic clusters during CSBs and iAPs. We found that the average cluster distance from the soma was significantly larger during CSBs than iAPs (Fig. 6E). These observations confirm earlier results suggesting the special importance of tuft branches in controlling CSBs in CA1 neurons (Takahashi and Magee, 2009; Bittner et al., 2015; Grienberger and Magee, 2022; Park et al., 2025).

To further analyze the synaptic and dendritic events leading to CSB firing, we turned to our extended currentscape method and calculated the average currentscapes for both CSBs and iAPs (Fig. 7A-B). The somatic currentscapes by current type and current origin revealed the strong Ca^2+^- and Na^+^-currents driving burst firing in the soma during CSBs and being responsible for the slight depolarization after iAPs. However, to our surprise, there was almost no difference between CSBs and iAPs until the end of the first spike neither in the magnitude, nor in the origin or the composition of the somatic currents.

**Fig. 7:**
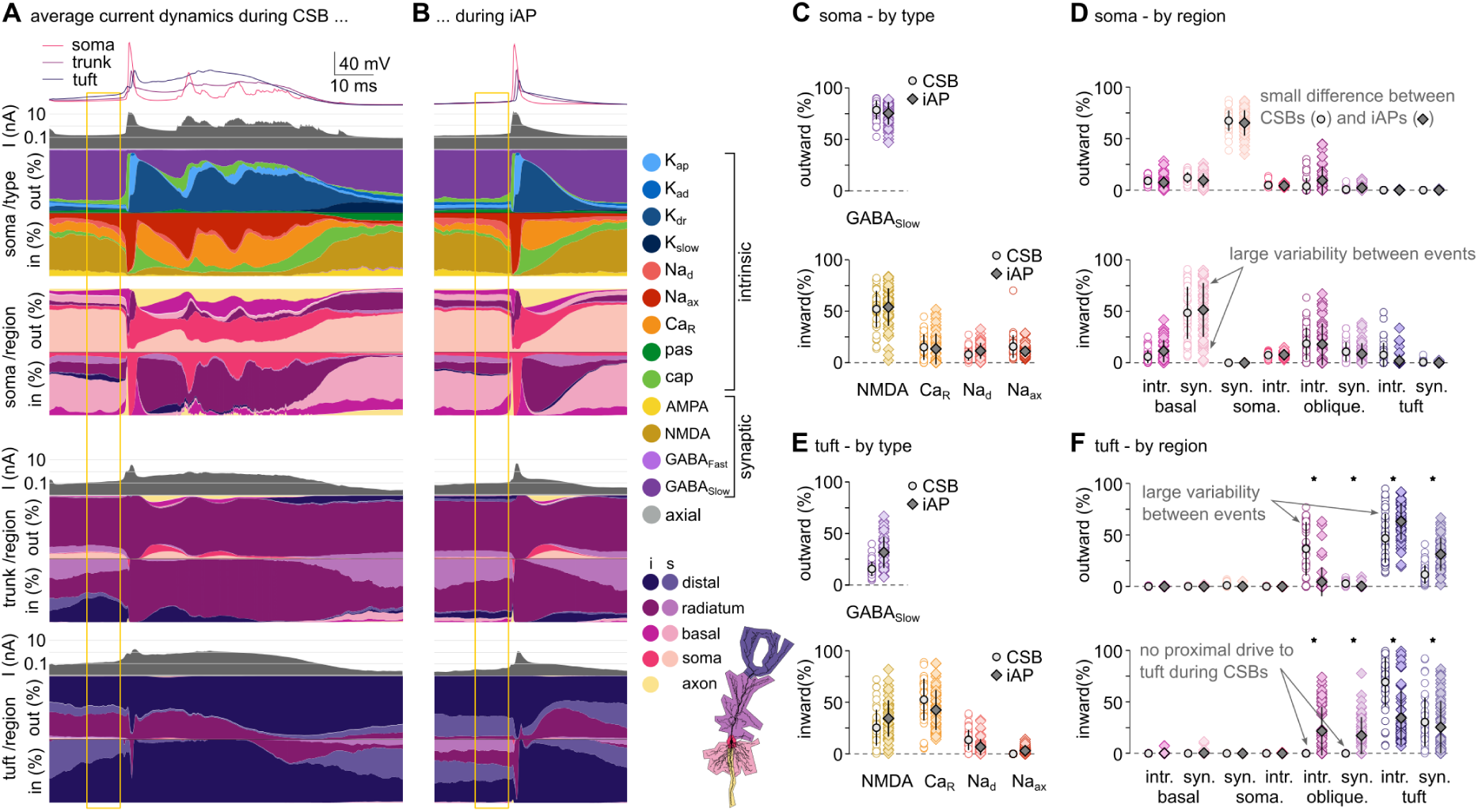
Currents underlying CSBs and isolated spikes. **A**) Average currentscape of CSBs, aligned to the first spike of the bursts (n=41 events). Top: average Vm in the soma, the trunk (d=260 *µ*m from soma) and in a tuft branch (d=470 *µ*m). Second row: Somatic total current and currentscapes by current type and current region. Third row: Total current and currentscape by current region in the trunk. Last row: Total current and currentscape by current region in the tuft. Yellow rectangle before the first spike indicates the region used for the analysis in C-F. **B**) Same as A for isolated spikes (n=58 events). **C**) Contribution of selected membrane current types to outward (top) or inward (bottom) somatic currents in individual CSBs (circles) or APs (dark diamonds). **D**) Contribution of different dendritic regions to the somatic currents in individual CSBs (circles) or APs (dark diamonds). **E-F**) Similar to C-D, calculated for a tuft dendrite.

Since we observed a large variability in the Vm, we speculated that averaging across many events could potentially conceal important differences between iAPs and CSBs. Therefore, we calculated the total magnitude of each current type in the soma for each event in the 8 ms period before iAPs or the first AP of CSBs (Fig. 7C). We found a large variability in the currents underlying both iAPs and CSBs with the distributions entirely overlapping between the two types of events. We got similar results when analysing the origin of the axial currents in the soma (Fig. 7D). This indicates that it is not possible to distinguish CSBs and iAPs before the first AP from the somatic state of the neuron, and suggests that the soma can not control burst firing. Consequently, the somatic current dynamics before the iAP and the CSB presented in Fig. 5Cc-Dd can be regarded as illustrative samples from a broad distribution, but the differences observed between them are not representative.

Next, we analyzed currents in the apical dendrites from the distal trunk region and from the apical tuft. Although we found some small, but significant alterations in the current types in both the trunk (not shown) and the tuft (Fig. 7E), the largest differences were in the origin of the currents driving these regions (Fig. 7F): before CSBs the tuft did not receive inward (excitatory) currents from proximal regions but was driven solely by local excitatory intrinsic and synaptic currents. In contrast, outward currents from the tuft propagated towards the trunk and obliques significantly more during CSBs than during iAPs. However, we still observed a substantial variability between the events indicating that both iAPs and CSBs could be highly diverse. We were wondering whether this diversity could be captured by a few dominant factors along which iAPs and CSBs were segregated.

In order to test this idea, we applied factor analysis to the dataset containing the magnitude of 42 current types in the 3 different dendritic domains (soma, trunk, tuft; see Methods) before the first spike of the event. The factor analysis revealed that the data indeed was low-dimensional: The first two factors together explained almost 80% of the total variance with additional factors providing negligible contribution (Fig. 8A). When we projected the data into the space defined by the first two factors, the datapoints corresponding to CSBs and iAPs tended to occupy different regions of the state space (Fig. 8B) indicating that these factors capture characteristic differences between these event types.

**Fig. 8:**
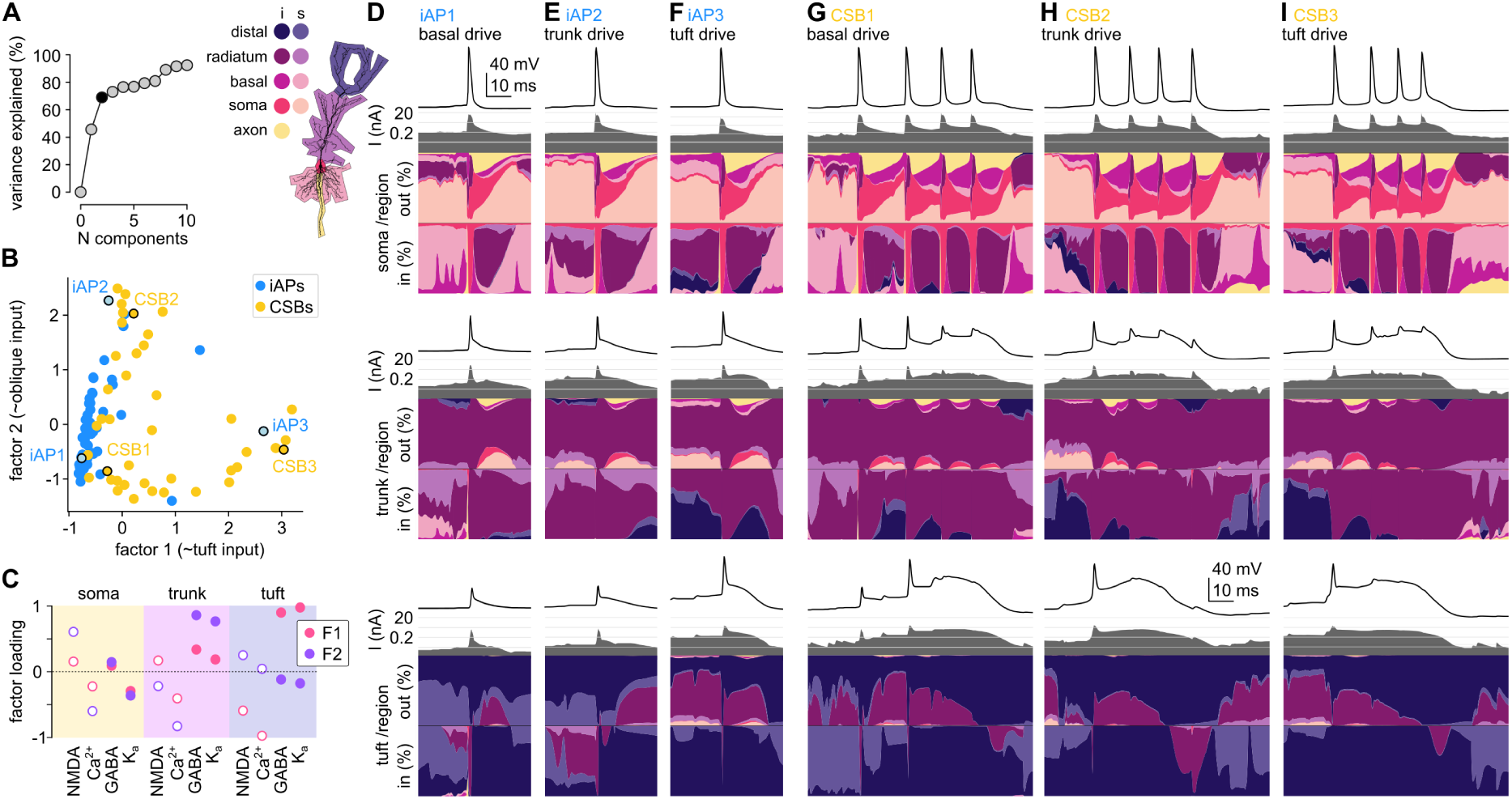
Structured diversity of the currents preceding CSBs and iAPs. **A**) Variance explained as a function of the number of components (factors) considered to reconstruct the magnitude of different current types preceding somatic events (CSBs or iAPs). **B**) Linear projection of the currents to the 2 factors explaining most of the co-variability across somatic events. The six events highlighted with black outline are shown in D-I. **C**) Factor loadings (weights) associated with representative inward (open circles) and outward (filled circles) currents of the two factors (F1: factor 1, pink; F2: factor 2, levander) shown in panel B. **D-I**) Example currentscapes of CSBs (D-F) and iAPs (G-I) with different initiation dynamics.

Next, we checked how different current types contributed to these first two factors. We found that the outward currents (GABA and K^+^) in the tuft region had the largest positive loadings, while the distal inward currents (NMDA and Ca^2+^ currents) had the largest negative loadings to the first factor (Fig. 8C; note that inward currents are negative). Thus, strong excitatory or inhibitory tuft inputs both increase the first factor. We thus interpreted the first factor as capturing the strength of the inputs to the tuft region. Similarly, the second factor had the strongest positive loadings on inhibitory, and negative weights on excitatory currents in the trunk region (Fig. 8C). Interestingly, proximal NMDA and Ca^2+^-current had opposite effects (Fig. 8C): somatic Ca^2+^-current, originating from the apical dendrites, increased both factors, while somatic NMDA currents, indicating dominant excitation from basal branches, decreased them. We interpret the second factor as capturing the strength of inputs to the oblique/trunk region of the cell.

This analysis revealed that most of the variability between iAPs is confined to a single dimension with low tuft input and a varying degree of oblique or basal excitation (Fig. 8B,D-E). Moreover, iAPs occur only exceptionally at high tuft input levels (Fig. 8F). In contrast, CSBs occupy a greater fraction of the state space with varying tuft and trunk input, but generally avoiding the lowest levels of tuft excitation (Fig. 8B, G-I). Note that occasionally highly similar currents preceded iAPs and CSBs and which of these occurred could depend on later synaptic inputs or subtle differences in the state of intrinsic conductances (Fig. 8B, D-I; see also Fig. S4).

To further elucidate the role of the inputs targeting different dendritic domains in evoking spikes or CSBs, we selectively decreased the input rates at basal, oblique or tuft branches, simulating optogenetic inhibition experiments (Bittner et al., 2015). Specifically, we randomly selected 300 input synapses targeting either of these domains, decreased their input rates by 75%, and measured the changes in postsynaptic spiking and CSB firing. We found that decreasing the tuft input had the strongest effect in reducing both the CSBs and the spike rate of the cell (Fig. S5A). This effect was not explained by the more clustered location of the inhibited synapses in the tuft (Fig. S5A), further emphasizing the crucial role of the tuft in controlling the excitability of the cell. In agreement with the experimental results (Bittner et al., 2015), the tuft inhibition caused a larger reduction in the CSBs rates (≈ 20% of the control) than in the spike rates (≈ 35% of the control; Fig. S5). Nevertheless, inhibition of basal or oblique synapses also decreased the number of CSBs, which confirms that complex spike bursts are not exclusively triggered by distal inputs in our model.

Taken together, these analyses demonstrated that, in agreement with previous experimental results (Takahashi and Magee, 2009; Bittner et al., 2015; Grienberger and Magee, 2022; Park et al., 2025) strong distal inputs facilitate the generation of CSBs. However, in our model there was no need for exceptionally high input synchrony at the tuft synapses neither for Ca^2+^-hotspots in the dendrites to evoke CSBs. Instead, CSBs could be evoked under a large variety of synaptic and intrinsic current conditions by input to apical dendrites even at relatively weaker tuft excitation.

## Discussion

### Summary

In this paper, we first introduced the extended currentscape technique to quantify and visualize the effect of dendritic events on the somatic response of a biophysical model neuron. Next, we illustrated the extended currentscape method in standard simulation experiments probing dendritic integration. Our method could compactly and intuitively capture the sequence of dendritic events leading somatic APs in models with simple morphology or to a superlinear somatic response in models with realistic dendritic arbor. After validating the model using spatiotemporally restricted stimuli, we turned towards *in vivo*-like inputs and examined dendritic activity during simulated place field dynamics. We found that although correlated activity dominated the dendritic Vm dynamics of the model, the extended currentscape method could identify various dendritic events underlying somatic spikelets, action potentials, and complex spike bursts.

Next, we contrasted the somatic and dendritic current dynamics preceding iAPs and CSBs. Although we found a large diversity of current contributions before the two event types, the distributions of the currents were surprisingly similar before CSBs and iAPs. Finally, we applied factor analysis to the somatic and dendritic currents and showed that the large variability is dominated by two factors, corresponding to inputs to the distal and proximal apical dendrites. We found that CSBs and iAPs could occur at variable oblique and basal input levels, but strong apical inputs facilitated the generation of CSBs.

### The extended currentscape

Our analysis is based on the extended currentscape method to identify the type and origin of membrane currents that drive the activity in any segment of a neuron with spatially extended morphology. Our technique relies on simulating the neuron using a standard biophysical modeling software (Hines and Carnevale, 1997) and measuring the membrane currents and the membrane potential in each model segment at every timestep. We used Kirchhoff’s current law to partition the axial currents between neighboring pairs of segments proportionally to the composition of the incoming or the outgoing membrane currents, depending on the direction of the axial current. We applied this partitioning algorithm iteratively starting from the tip of the dendrites (leaf nodes) and proceeding towards the target segment.

In this paper, we partitioned the axial currents either by the channel type or by the dendritic region of the associated membrane currents. Partitioning the axial currents by the type of the membrane currents generating them provides a compact and intuitive summary of the sequence of events mediated by different intrinsic and synaptic conductances during various neuronal activity patterns, illustrated here for the case of theta activity, spiking and bursting in place cells. In contrast, partitioning the axial currents by their dendritic origin reveals the relative contribution of the different dendritic domains to driving the neural response at any given time. Here we distinguished only a few major regions (axon, soma, basal, oblique and tuft dendrites) within the cell, but different divisions, with more fine grain compartmentalization or alternative combinations with partitioning by current type, would be an interesting future application of the method.

Previous approaches dissecting the contribution of individual ion channel types to the somatic response of a cell relied on targeted perturbation of the available biophysical mechanisms both in real neurons (Wei et al., 2001; Ariav et al., 2003; Losonczy and Magee, 2006; Smith et al., 2013; Palmer et al., 2014) and in biophysical models (Gasparini et al., 2004; Gómez González et al., 2011; Hay et al., 2011; Behabadi et al., 2012; Grienberger et al., 2014; Goetz et al., 2021; Kim et al., 2023). However, interpreting the result of these perturbations is hampered by the potential interactions between various biophysical mechanisms, especially under *in vivo* conditions (Smith et al., 2013; Palmer et al., 2014). For example, Losonczy and Magee (2006) found that blocking dendritic Na^+^-spikes inhibited both the fast and the slow components of the superlinear somatic response. They posited that the supralinear input-output function critically depends on the facilitation of NMDA spikes by local dendritic Na^+^spikes. Conversely, Smith et al. (2013) found the blocking NMDA receptors greatly reduced the number of local dendritic fast spikes in layer 2/3 PNs, suggesting a reciprocal interaction between NMDA– and Na^+^–spikes. Here we propose an alternative approach that does not require perturbations but calculates the contribution of ion channel types directly from the recorded axial and membrane currents, albeit currently only in biophysical models. Importantly, our method works equally well under a wide variety of input conditions, including stimulation localised to a single dendritic branch or distributed throughout the entire dendritic tree.

### Biophysical model

Our biophysical model CA1 neuron is a modified version of the model of Jarsky et al. (2005) originally developed to capture the generation and propagation of Na^+^ spikes along the apical trunk and later modified to account for the integration of synaptic inputs via Na^+^-, and NMDA-spikes in basal and oblique dendrites (Losonczy and Magee, 2006; Ujfalussy and Makara, 2020). As Ca^2+^-spikes in CA1 PNs are primarily mediated by R-type channels (Takahashi and Magee, 2009), we added Ca^2+^channels using kinetic scheme taken from (Magee and Johnston, 1995) to model dendritic Ca^2+^spikes. Our model was able to generate complex spike bursts, with associated dendritic plateau potentials upon current injection or synaptic stimulation into the dendrites. However, this model is not yet able to capture all, potentially important, properties of Ca^2+^-spikes in CA1 neurons, including their failure to generate Ca^2+^-spike upon strong somatic current injection in most cells, the delayed initiation of the Ca^2+^spikes, the inhibitory effects of the Na^+^-APs on Ca^2+^ spike initiation (Golding et al., 1999) and the generation of prolonged somatic plateau potentials (Bittner et al., 2015). Currently we are not aware of any CA1 PN model that would be able to reproduce these experimental observations (see the references in Sáray et al. 2021) indicating the need for further research revealing the exact biophysical mechanisms of dendritic Ca^2+^spikes.

The membrane potential response of the model to *in vivo*-like synaptic inputs was dominated by a low-dimensional, global dynamics (Fig. 4C-E). The shape and the variance explained by the first two principal components of Vm covariance matrix was very similar to the data reported in a recent *in vivo* voltage imaging study (Lee et al., 2026), indicating that back-propagation of APs is the most prominent event organising dendritic activity. However, our model also exhibited a lot of local dendritic events each restricted to only a few branches (Fig. 4H-J). Importantly, in our model localised dendritic events occurred primarily in distal, terminal branches that are more likely to be omitted during single-cell voltage imaging studies (Lee et al., 2026). In our model the proximal dendrites were highly synchronized with the soma, whereas previous Ca^2+^-imaging studies reported some level of uncoupling of somatic and dendritic activity (Sheffield and Dombeck, 2015; Sheffield et al., 2017; Rolotti et al., 2022). This may be explained by the unreliability of Ca signal measurements to report Vm changes (Tran-Van-Minh et al., 2016; Weber et al., 2016; Landau et al., 2022). Our analysis also indicates that global events may dominate the PCA analysis (Lee et al., 2026) even when local dendritic regenerative events are widespread, and the neuron is mainly driven by strong, functionally clustered synaptic inputs to a few dendritic branches (Ujfalussy and Makara, 2020).

We used identical biophysical properties and input statistics for the tuft and distal oblique branches in our model. Although numerous differences are known regarding both the inputs (Witter et al., 1989; Isomura et al., 2006; Milstein et al., 2015) and dendritic physiology (Nicholson et al., 2006; Bittner et al., 2012), many of the relevant parameters are still underconstrained. We wanted to avoid overfitting by keeping the model relatively simple (Herz et al., 2006). Moreover, by maintaining identical inputs and ion channel distribution we could distinctly highlight the special role of tuft morphology in CSB generation and altering the inputs or the ion channel density for the tuft would make these interpretation of the results more ambiguous. Elucidating the specific role of the regional differences in inputs or their local processing in CSB generation could be the subject of future investigations.

### Implications for burst firing

While acknowledging the incompleteness of our biophysical model, we believe that our simulations still provide several important insights into the generation of Ca^2+^-spikes under *in vivo*-like conditions. First, it was unclear whether increased Ca^2+^-channel density in the tuft region or special perforant path input patterns are necessary for facilitating burst firing in CA1 PNs (Bittner et al., 2015; Grienberger and Magee, 2022). Our simulations demonstrated that distal tuft branches are especially well suited to promote burst firing even with uniform Ca^2+^channel density and similar input statistics along the apical subtree. This observation suggests that it is the morphology of the cell and the presence of strong inhibitory conductance load on the apical trunk that makes tuft branches more excitable than oblique dendrites.

Second, we observed a high variability between the activation of individual tuft and oblique branches during CSB firing. This observation is reminiscent of Ca^2+^ imaging data from the distal dendrites showing a variable recruitment of tuft branches during place field formation and subsequent traversals (O’Hare et al., 2025). Our simulations clarified that tuft Ca^2+^-plateaus are not necessarily associated with uniform tuft activation and the high variability between branches can be caused by diversity in the synaptic inputs they receive.

Third, we found a large heterogeneity among the input currents preceding CSBs: while some CSBs were elicited at high tuft currents, many were triggered at lower or intermediate level of distal inputs. This suggests that the presence of distal inputs facilitates rather than controls the emergence of CSBs. This finding explains why it is possible to elicit plateau potentials *in vivo* using somatic current injection during BTSP (Behavioral Time Scale Plasticity) induction (Bittner et al., 2015) where the same stimulus usually fails to evoke Ca^2+^-spikes *in vitro* (Golding et al., 1999) and predicts that reduction of CA3 inputs to the oblique or basal dendrites would also decrease CSB rate, BTSP induction and the associated reward zone over-representation similarly to the effect of inhibiting the more distal EC inputs (Fig. S5; Bittner et al. 2015; Grienberger and Magee 2022; Fan et al. 2023). The facilitatory action of distal tuft inputs on CSBs is reminiscent of the conjunctive bursting mode, where bursts are generated by a synergistic interaction between different input streams (Larkum et al., 1999; Naud and Sprekeler, 2018; Takahashi and Magee, 2009) and is consistent with the conclusions of the recent study postulating that dendritic plateaus are initiated within the distal regions of stratum radiatum by strong inputs to both distal tuft and radial oblique dendrites (O’Hare et al., 2025). The ability of CA3 inputs to the radial oblique dendrites to influence CSB generation and BTSP induction could be an important cellular mechanism to enable generalization across consecutive experiences (Vaidya et al., 2025; Qian et al., 2025).

Although we could frequently observe complex spike bursts in our simulations, we did not observe large amplitude (>20 mV) prolonged (> 100 ms), plateau-like depolarization events in the soma with substantially reduced AP amplitude (Bittner et al., 2015) during naturalistic synaptic inputs. However, a response more similar to plateau potentials could be evoked in our model by combining *in vivo*-like synaptic stimulation with direct somatic current injection (Fig. S1E). We thus speculate that the generation of plateau potentials might require strong perisomatic excitatory currents. Our preliminary simulations suggest that Ca^2+^-channels added to the basal dendrites can provide this additional excitation. The biophysical mechanism and the natural synaptic input conditions that lead to CSB versus plateau potentials in CA1 pyramidal neurons is a promising subject for future research.

### Limitations of the study

In order to make the partitioning algorithm self-consistent, we had to also include the capacitive current among the membrane currents in Eq. (1): It is possible that in a given segment the only inward current may be the capacitive current but the axial current is still outward. The capacitive current acts as a delay line in the membrane equation, representing the indirect effect of currents that were active earlier and had charged the membrane capacitance. In practice, we found that the capacitive current may sometimes have a large contribution to the membrane dynamics (Fig. 5Ce) and can thus mask the delayed contribution of various membrane currents. A potential extension of our method would be to also partition the capacitive current, but it is not clear how this could be achieved self-consistently and is beyond the scope of our current paper.

Our partitioning algorithm identifies only the direct contribution of the membrane currents to membrane potential changes in the target compartments, and ignores all indirect effects. This property follows from the presumption that there must be a direct chain of links between cause and effect (Pearl, 2009), in particular, there must be a continuous flow of axial current between the membrane current in a distal dendrite and the change it triggers in the target. Thus, our method is blind to indirect effects otherwise known to be present in complex dendritic trees, such as certain types of off-the-path inhibition (Gidon and Segev 2012; but see also Fig. 2P). Revealing the contribution of such indirect causes under complex synaptic stimulation would require causal perturbation methods that have been proposed earlier in the context of synaptic effects (Bicknell and Häusser, 2021).

### Testing the model: electrophysiology or voltage imaging

Experimental testing of the partitioning algorithm presents considerable challenges. Accurate measurements of the contribution of membrane current types to the somatic activity of a neuron require simultaneous measurement of all membrane currents and potentials, which is not feasible with current experimental methods.

Although measuring individual current contributions is not feasible in real neurons, a recent study performed whole-cell recordings in visual cortical neurons *in vivo* and used the systematic change of the input resistance with depolarization to estimate the contribution of intrinsic and synaptic currents to neuronal responses (Li et al., 2020). They found that during visual stimulation the intrinsic and synaptic conductances have comparable contribution to the subthreshold membrane potential changes of the cell, with intrinsic channels amplifying the synaptic response. Similar analysis on *in vivo* hippocampal recordings could test whether synaptic and intrinsic conductances amplify or counteract each other *in vivo* (Hoffman et al., 1997; Bittner et al., 2015; Epsztein et al., 2011; Cohen et al., 2017; Valero et al., 2022). However, in the absence of dendritic recordings, this study was unable to identify the spatial origin of synaptic and intrinsic changes within the dendritic tree.

More direct test of current propagation in real neurons would require measuring the membrane and axial currents at multiple spatial locations. As a recent step towards achieving this Meszéna et al. (2023) combined somatic patch-clamp recordings and multichannel extracellular recordings to reconstruct the spatiotemporal distribution of the membrane currents and the membrane potential of a single neuron during AP generation. Alternatively, one could use *in vivo* voltage imaging to monitor the membrane potential of dendrites and the soma simultaneously (Abdelfattah et al., 2023; Liao et al., 2024; Park et al., 2025; Lee et al., 2026). From the local membrane potential one can calculate the axial currents after estimating the intracellular resistivity and the dendritic cable diameters. Our method of partitioning the axial current by its origin within the dendritic tree can be applied directly to this kind of data. Therefore, such data could be used to directly test both the behaviour of the biophysical model under *in vivo*-like input conditions (Fig. 4F-G) and the diversity in the origin of the input currents before single spikes and CSBs in CA1 neurons (Fig. 8).

## Methods

### Biophysical models

All simulations were performed with the NEURON simulation environment (version 8.2) embedded in Python (version 3.9 and 3.11). Code for simulating the biophysical model, preprocessing, axial currentscape calculation and visualization are available at https://github.com/bencefogel/currentscapes-invivo-demo. Code for simulating the simplified model is available at https://github.com/bencefogel/currentscapes-simple-model-examples.

### Simplified neuron model (**Fig. 2**)

A simplified model was constructed using a minimal morphology consisting of a soma and three dendritic sections. The soma was connected to a primary dendrite (dend1), which bifurcated into two secondary dendrites (dend2 and dend3). All sections were modeled as unbranched cylinders.

The soma had a diameter and length of 20 *µ*m. The primary dendrite (dend1) was 100 *µ*m long with a diameter of 2 *µ*m and was discretized into 11 segments to ensure adequate spatial resolution. The two secondary dendrites (dend2 and dend3) each had a length of 50 *µ*m and a diameter of 1.5 *µ*m, with 5 segments per branch. Dend2 and dend3 were attached to the distal end of dend1, forming a simple three-branch dendritic arbor.

Specific membrane capacitance was set to 1 *µ*F/cm^2^. Passive leak conductance was implemented using NEURON’s pas mechanism, with a uniform leak reversal potential of -66 mV. The soma had a reduced leak conductance (1/40000 S/cm^2^), whereas dendritic sections used a leak conductance of 1/20000 S/cm^2^. Axial resistivity (Ra) was set to 100 Ωcm for the soma and 800 Ωcm for all dendritic compartments.

Active voltage-gated currents were included in the soma and dendrites to support basic excitability. The kinetic parameters of the active conductances were identical to those in the morphologically detailed model (see below; Fig. S6). The soma contained fast sodium and delayed-rectifier potassium channels with maximal conductances of *g*_Na_ = 0.02 S*/*cm^2^ and *g*_K_ = 0.002 S*/*cm^2^, respectively. All dendritic compartments expressed lower-density dendritic sodium and delayed-rectifier potassium currents with *g*_Na,dend_ = 0.007 S*/*cm^2^ and *g*_K,dend_ = 0.0002 S*/*cm^2^, providing modest active boosting and limited back-propagation. Synaptic mechanisms included AMPA and GABA_fast_ receptors implemented using Exp2Syn. AMPA synapses used *τ*_1_ = 0.1 ms and *τ*_2_ = 1 ms with a reversal potential *E*_AMPA_ = 0 mV, while GABA_fast_ synapses used *τ*_1_ = 0.1 ms, *τ*_2_ = 4 ms, and *E*_GABA_ = −65 mV with a synaptic conductance *g*_GABA_ = 10 nS. Excitatory synaptic strengths varied by experiment: AMPA conductance was set to 0.5 nS for Fig. 2A-C and O, and 2.5 nS for Fig. 2N and P.

Synaptic inputs were generated using NetStim objects in NEURON to produce brief, high-frequency bursts of presynaptic events. Each stimulus (for both AMPA and GABA) consisted of five spikes delivered at 0.1 ms intervals, corresponding to a set of synchronous presynaptic spikes designed to evoke localized dendritic responses.

### CA1 neuron model

We used a modified version of a previously developed CA1 PN model (Jarsky et al., 2005; Ujfalussy and Makara, 2020), maintaining the same ion channel distributions and densities for Na^+^, K^+^ and proximal and distal type K^+^ channels as described in Ujfalussy and Makara (2020) (Fig. S6).

To capture dendritic Ca^2+^ spikes and somatic CSBs, we added an R-type Ca^2+^ channel to the model. The Ca^2+^ channel kinetics were based on the steady-state activation and inactivation curves fitted by Magee and Johnston (1995), with a slight modification to allow for a larger window current (Fig. S1A). The Ca^2+^ channel was expressed in the apical dendritic tree with a uniform distribution and an ion channel density of 0.006 S*/*cm^2^.

Furthermore, we added a high-voltage activated, slow K^+^ channel that was used to simulate the combined effect of multiple K^+^ channels including Ca^2+^-activated K^+^ channels, contributing to the repolarization after the Ca^2+^ spikes. The slow K^+^ channel was expressed in the apical dendritic tree with a uniform distribution and an ion channel density of 0.001 S*/*cm^2^.

The idealized CA1 place cell received inputs from 2000 excitatory and 200 inhibitory presynaptic neurons. We simulate the activity of excitatory and inhibitory neurons during the movement of a mouse on a 2-m long circular track with constant velocity of 20 cm/s. Each presynaptic excitatory neuron had a single idealised place field and represented the CA3 inputs received by the postsynaptic CA1 neuron.

Reliable place cell tuning can be achieved by functional synaptic clustering without increased excitatory drive in the place field (Ujfalussy and Makara, 2020) or via strong excitatory drive without input clustering (Grienberger et al., 2017; Ujfalussy and Makara, 2020). However, experimental data indicates that both of these mechanisms are present and contribute to the activity of place cells (Adoff et al., 2021; Tasciotti et al., 2025). Excitatory synapses were placed randomly with a uniform distribution on the entire dendritic tree, except 240 inputs, active in the postsynaptic place field in middle of the track, that were selected to form functional synaptic clusters. There were a total of 12 clusters, and each cluster had 20 synapses. Synaptic clusters were placed on the middle of terminal dendritic branches longer than 60 *µ*m with 1 *µ*m distance between the synapses. This relatively strong clustering favoured the generation of local dendritic spikes (Ujfalussy and Makara, 2020).

Although interneurons can display spatial tuning, they typically have a broad tuning with low selectivity (Ego-Stengel and Wilson, 2007; Dupret et al., 2013; Geiller et al., 2020). Albeit weak disinhibition within the place field can contribute to the selective firing of place cells (Geiller et al., 2022; Valero et al., 2022), this was not necessary for place cell activity in novel environments (Geiller et al., 2022) and the overall inhibitory input to place cells is largely untuned (Grienberger et al., 2017). Inhibitory presynaptic inputs were modulated by theta oscillation (Csicsvári et al., 1999) but were not spatially tuned (Wilson and McNaughton, 1994; Dupret et al., 2013). The inhibitory synapses were divided into two groups with 80 synapses targeting the soma and the apical trunk and the remaining synapses distributed randomly along the entire dendritic tree (Megias et al., 2001).

The model included AMPA and NMDA excitation and slow and fast GABA inhibition. Synaptic parameters were kept identical to those described in Ujfalussy and Makara (2020) (see Fig. S7 for kinetic parameters). To induce reliable firing within the neuron’s place field, we increased the AMPA (from *g*_max_ = 0.6 *nS* to *g*_max_ = 1 nS) and NMDA (from *g*_max_ = 0.8 nS to *g*_max_ = 1.2 nS) conductance associated with the clustered synaptic inputs, consistent with the primary role of synaptic plasticity in place field formation (Bittner et al., 2015; Adoff et al., 2021; Heredi et al., 2025). Inhibitory synapses (both slow and fast) had a maximal conductance of *g*_max_ = 0.2 nS.

### Simulation of the model

We simulated place cell activity for 10 seconds as the hypothetical animal completed a single lap on the track. A total of 16 simulations were run, each with different random configurations of synapse placement and presynaptic input patterns.

In the NEURON software, neurons are modelled as a series of sections, which represent unbranched parts of continuous cable, such as dendrites or axon. These sections are connected together to form an acyclic graph, according to the morphology of the neuron. Each section is divided into multiple segments of equal length.

NEURON uses a normalized distance to express locations along a section, where 0 represents the start (closest to the parent section) and 1 represents the distal end. To compute membrane dynamics, NEURON calculates membrane potentials and membrane currents at discrete positions, known as internal nodes, located at the center of the segments. The accuracy of the partitioning, like the accuracy of the numerical simulation of spatially extended cables, can be increased by choosing finer discretization, ie., larger number of segments. During our simulations, the number of segments within each section was chosen to ensure accurate description of signal propagation within the cable with reasonable computational efficiency. As a rule of thumb, we found that axial current partitioning is accurate if the proportion of the magnitude of the membrane currents and the magnitude of the axial current was small. This proportion was on average ≈ 1 : 17 for non-terminal branches in our case.

In addition to the internal nodes, located at the center of the segments, there are external nodes located at the 0 and 1 ends of each section. These external nodes are only used to connect segments, but no membrane mechanisms (synapses or ion channels) are associated with these external nodes.

During each simulation, we recorded the membrane potential (mV), intrinsic (mA/cm^2^) and synaptic currents (nA) of all segments of the neuron (*N*_sec_ = 161 sections and *N*_seg_ = 1529 segments in total, including the 2*N*_sec_ internal segments; see below). This was the raw dataset we used to calculate the currentscapes. We also saved the model connectivity structure between segments together with the axial resistance values (MΩ) to calculate axial currents offline. Finally, we measured the area (*µ*m^2^) of each segment, which is needed during the preprocessing of membrane currents.

During the simulation, state variables were computed using NEURON’s built-in multi order variable time-step integration method. For subsequent preprocessing and currentscape calculation, the recorded output vectors were downsampled to 5 kHz.

### Preprocessing

The goal of preprocessing was to convert the raw dataset, saved by NEURON into two tables that we can use to calculate the partitioning of the axial currents. The first table contains the membrane currents for each segment and for each time step. The second table contains the axial currents between the segment pairs.

### Preprocessing membrane and synaptic currents

The preprocessing starts with choosing a target section for partitioning. The extended currentscape will partition the axial currents flowing into this target section. If we want to partition the axial currents flowing into different sections (e.g., soma and a tuft compartment), we have to repeat the pruning (see below) and the partitioning steps with the novel target.

All segments that belong to the target section were merged for further analysis (that is, the membrane currents of the segments were summed and no axial current was calculated within the target section). The connections between the segments of the target section and its child branches were reassigned to the segment of the merged target.

Each synapse is represented by a conductance-based mechanism that injects current into the post-synaptic compartment when triggered via a NetCon object. Since each NetCon delivers spikes independently, and multiple synapses can target the same segment, synaptic currents (in nA) were summed per segment and per synaptic type (AMPA, NMDA, GABA_fast_, GABA_slow_) to accurately capture the total synaptic input.

To ensure consistency in subsequent calculations, intrinsic currents were converted from units of mA/cm^2^ to nA. This was done by multiplying the recorded current by the corresponding segment area (in *µ*m^2^) and applying a scaling factor of 10*^−^*^2^. Finally, intrinsic and synaptic currents were combined into a single data structure, where each row corresponds to a unique segment and current type pair, and each column is a simulation time point.

The core algorithm underlying the extended currentscape calculation is the axial current partitioning algorithm (see below). By default, this algorithm decomposes the axial current of a target compartment into underlying membrane current components. However, alternative variants of the partitioning algorithm can be applied if the membrane current data structure is reindexed and recalculated. In region-specific partitioning, membrane currents are reassigned based on their location within different subcellular regions (for example, axon or dendrite). These different approaches can also be combined in a flexible way to measure e.g., the somatic influence of NMDA channels in a particular dendritic branch. In this study, we demonstrate two such variants: Membrane current type-specific partitioning and a combined partitioning where we distinguished 5 subcellular regions (axon, soma, basal dendrites, trunk+oblique dendrites and tuft dendrites) and 2 current types (synaptic and intrinsic currents). The intrinsic category incorporated all non-synaptic currents, including capacitive and passive currents.

### Preprocessing axial currents

To compute axial currents we first created a list with the adjacent segment pairs between which axial currents will be calculated. Connections between sections (branches) are implemented in NEURON through a pair of external nodes: one at the distal end of the parent section and another at the proximal end of the child section. Although these nodes formally belong to different sections (parent and child), they have identical membrane potential as they represent the same point within the cable, so we merged them to remove the duplicates from our primary dataset.

The axial current between a parent and child segment is computed using Ohm’s law:

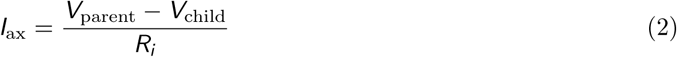

Here *V*_parent_ and *V*_child_ denote the membrane potentials of the parent and child segments, respectively, and *R_i_*denotes the axial resistance between them.

### Extended Currentscape Calculation

The extended currentscape calculation requires the following inputs: the preprocessed membrane and axial currents across all segments, a specified target segment, and a defined partitioning strategy, such as current type-specific or region-specific partitioning.

The first step in calculating the extended currentscape was to separate positive and negative membrane currents. Certain currents, such as capacitive currents, can be both inward and outward at the same time in different compartments. Without separation, such positive and negative currents would partially cancel each other out, reducing the contribution of the current type during the extended currentscape calculation. This effect is even more pronounced when partitioning the axial currents by the origin (see below): Strong positive and negative currents from two different basal branches can cancel each other, largely concealing their overall contribution to the somatic activity. By treating the positive and negative membrane currents independently, we ensured that the outward and inward membrane current components of the target’s axial current are calculated separately.

Following preprocessing, including reindexing and recalculation of membrane currents when required by the region-specific partitioning, a directed graph was constructed at each time step. The nodes of the graph represent individual neuronal segments and the directed edges indicate the direction of axial currents flowing between them. We create two separate subgraphs starting from the target segment: one for the inwardly flowing and one for the outward axial currents. Partitioning is calculated independently for the two subgraphs, with the inward (outward) axial graph partitioned according to the inward (outward) membrane currents, respectively.

To determine the partitioning order, we performed a depth-first search (DFS) on the two directed graphs starting from the target segment mowing towards the terminals along each dendritic branch separately. The search proceeds through the directed graph until it encounters collision points, where the axial current reverses its direction. At these points, current propagation is blocked: Distal nodes beyond the the first reversed edge (*collision edge*) are excluded from further calculations, as they do not directly influence the target compartment. In a benchmark simulation this pruning process reduced the size of the tree (which is proportional to the computational cost of the currentscape calculation) by 68% (50–80%). The two subgraphs identified by this search define the structures upon which axial current partitioning is performed.

Partitioning is carried out iteratively, beginning from the leaf nodes of the subgraph and progressing toward the target node. At each time step, the axial current is partitioned into inward (negative) or outward (positive) membrane current components depending on the direction of the axial current flow. Specifically, for a given time point *t*, the positive or negative membrane currents *I* ^m,child^ of the child node are first normalised by their sum. The resulting fractions are multiplied by the magnitude of the axial current *I* ^ax^ flowing between the child and the parent nodes. The contribution of membrane current of type *i* to the axial current can be expressed as:

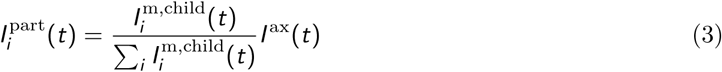

The partitioned axial current components *I* ^part^(*t*) are then added to the corresponding membrane currents of the parent section according to:

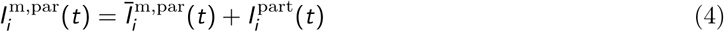

where 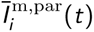 is magnitude of the membrane current type *i* measured in the parent segment. This procedure is repeated iteratively for each pair of nodes along the subgraph until the target compartment is reached.

The final output of the extended currentscape calculation consists of two data structures, each including the partitioned axial currents of the target compartment: one containing the inward membrane currents and the other containing the outward membrane currents in each time step. Simulating the morphologically detailed CA1PN model and performing the partitioning at 5 kHz temporal resolution on a 1 s long simulation took 75 min. on a MacBook Pro (2019) computer with 2.6 GHz 6-Core Intel Core i7 processor and 16GB memory.

## Data Analysis

### CSBs and isolated spikes

CSBs were detected by first smoothing the somatic Vm with a 20 ms Gaussian kernel and then detecting the crossing of a -55 mV threshold of the smoothed subthreshold Vm (Fig. 4B). The start of the CSB was defined as the time of the first spike after crossing this threshold. Isolated spikes were defined as action potentials with a distance of at least 30 ms from the closest AP.

### Ca^2+^-spike detection

Dendritic Ca^2+^-spikes (Fig. 6C-D) were detected by smoothing the dendritic Vm with a 4 ms Gaussian kernel and then detecting the crossing of a -35 mV threshold of the smoothed subthreshold Vm (Bittner et al., 2015).

### PCA of dendritic voltage

We recorded the Vm in the middle segment of all dendritic and somatic compartments of the model and calculated the covariance matrix of the z-scored dendritic voltages in each run separately. Fig. 4C shows the average of these 16 covariance matrices. PCA was then performed by calculating the eigenvalues and eigenvectors of the covariance matrices separately (Fig. 4D-E).

### Dendritic events

Dendritic Vm peaks were detected in the central segment of each dendritic section by the find_peaks function from the signal modul of the scipy python package with parameters height=-30 mV and prominence=30 mV. Fig. 4F shows the histogram calculated from the time difference between dendritic events in all branches and the closest somatic action potential. We defined the spatial extent of dendritic events as the number of interconnected dendritic branches that each showed a local Vm peak (as defined above) within *<* 2 ms of the Vm peak in at least one of the neighbouring branches.

### Dendritic domains

In Fig. 6A-B we show the average Vm in different dendritic domains during CSBs and isolated APs. We included all terminal branches with *L >* 60 *µm* in this analysis (basals: *N*_basal_ = 30, average distance from the soma: *D*_basal_ = 135 *µm*; obliques: *N*_oblique_ = 19, *D*_oblique_ = 313 *µm*; *N*_tuft_ = 9, *D*_tuft_ = 729 *µm*;).

### Average currentscapes

To calculate the average currentscapes in Fig. 6 we first calculated the percentage of the different current shares of each event and then averaged the percentages across the events.

### Factor analysis

Factor analysis in Fig. 8 was performed using the partitioned somatic, trunk, and tuft current types before the first spike of the CSB or iAP events. Specifically, we averaged the magnitude of the incoming current types in the interval *t* = {−10, −2} before each event. We performed the averaging separately for the positive and negative currents for all types with nonzero variance (e.g., NMDA currents are always inward, so their contribution to outward currents has zero mean and variance). For this analysis, we used the raw current magnitudes instead of the percentages. We performed factor analysis using the FactorAnalysis function using sklearn’s decomposition module.

### Simulating optogenetic inhibition

We simulated targeted optogenetic inhibition of different dendritic domains (Fig. S5) by randomly selecting a similar number (E[*n*] = 300) of presynaptic inputs and decreasing their firing rate to 25% of their baseline value. In different experiments, we selected inputs targeting either basal, oblique, or tuft dendrites.

## Acknowledgements

We thank Judit K Makara, Szabolcs Káli and János Brunner and all members of the Neuronal Signalling and the Biological Computation Research Groups for discussions and for their comments on the manuscript. B.B.U. was supported by the National Research, Development and Innovation Office of Hungary (FK-125324) and by the Hungarian Research Network.

## Supplemental Figures

**Fig. S1:**
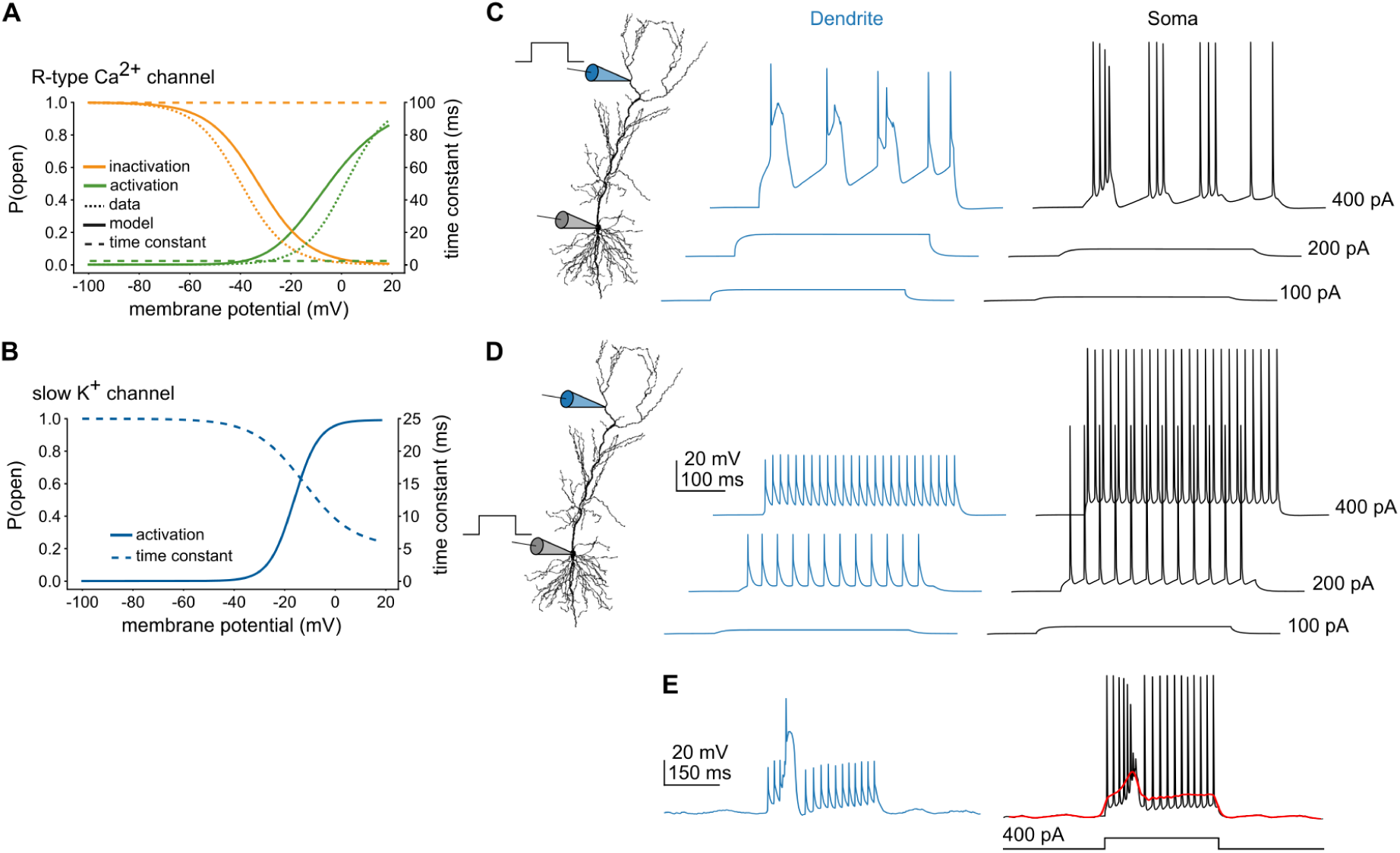
Biophysical model of burst firing in CA1 PNs. **A**) Steady-state activation and inactivation curves (left axis, solid lines) and the time constants (right axis, dotted) of the R-type Ca^2+^channel used in the model. Dotted lines illustrate sigmoid curves fitted to data in Magee and Johnston (1995). **B**) Steady-state activation curve (left axis, solid) and time constant (right axis, dashed) of a slowly activating K^+^ channel. **C-D**) Dendritic (blue) and somatic (black) Vm response of the model to dendritic (C) and somatic (D) current injections. The model has a lower current-threshold for Na^+^-spikes in the soma and for Ca^2+^-spikes in the dendrites. Note, that CSBs can be triggered by stronger somatic current steps in the model (not shown). For similar experimental data see Golding et al. (1999). **E**: Dendritic and somatic Vm responses of the model to a 300 ms, 400 pA somatic current injection under in vivo-like synaptic input conditions (during theta activity, outside the place field, as in Fig. 5B). The red line indicates the smoothed somatic Vm. Under these conditions, a dendritic Ca^2+^-spike and an associated somatic CSB can be evoked by somatic current injection. For similar experimental data see Bittner et al. (2015).

**Fig. S2:**
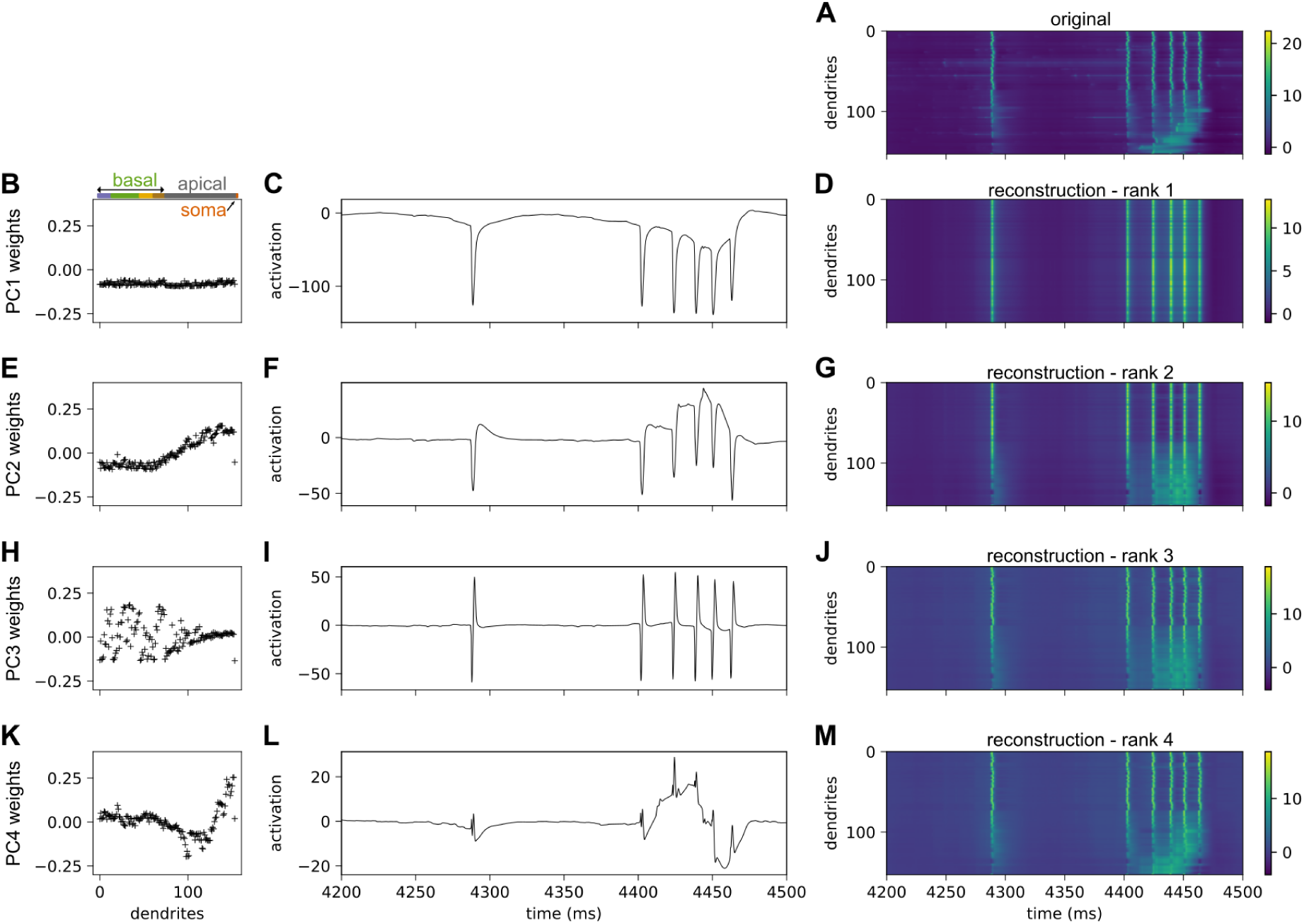
Dynamics of dendritic Vm PCA components. **A**) Short segment of the z-scored Vm of the dendritic branches (rows; soma is the last) of the simulated neuron during *in vivo*-like synaptic stimulation. The cell fired an action potential around *t* = 4300 ms and a CSB around *t* = 4400. Dendrites are ordered by morphology as in Fig. 4E. **B)** The weights of the first component (PC1) in an example simulation. Dendritic branches are ordered as in Fig. 4E. **C)** The temporal activation dynamics of the first component (PC1) in an example simulation on the same segment shown in A. **D**) Reconstruction of the dendritic membrane potentials using the first PCA components. **E-M** Similar to A-C for the second (E-G), third (H-J) and fourth (K-M) PCA components. The reconstructions in panels G, J and M used all components up to the given rank. Related to Fig. 4

**Fig. S3:**
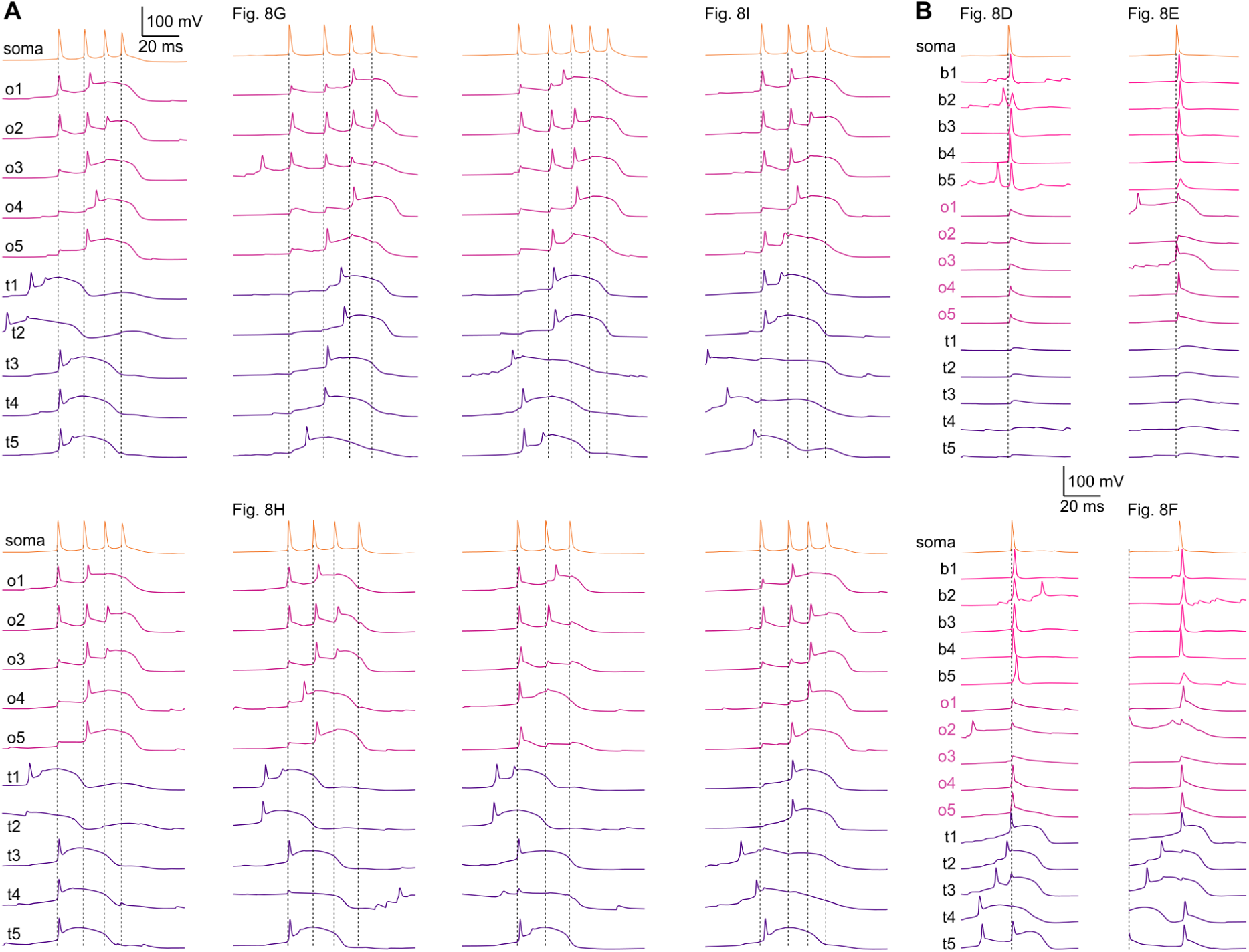
Further examples of somatic and dendritic Vm dynamics during CSBs and iAPs. **A**) Vm response of 5 oblique and 5 tuft branches during 8 different CSB events. The recording locations are the same as in Fig. 4F. The events shown in Fig. 8G-H are indicated above the traces. The timing of the somatic action potentials are marked by dotted vertical lines. **B**) Vm response of 5 oblique and 5 tuft and 5 basal branches during 4 different iAP events. The events shown in Fig. 8D-E are indicated above the traces. Related to Fig. 4.

**Fig. S4:**
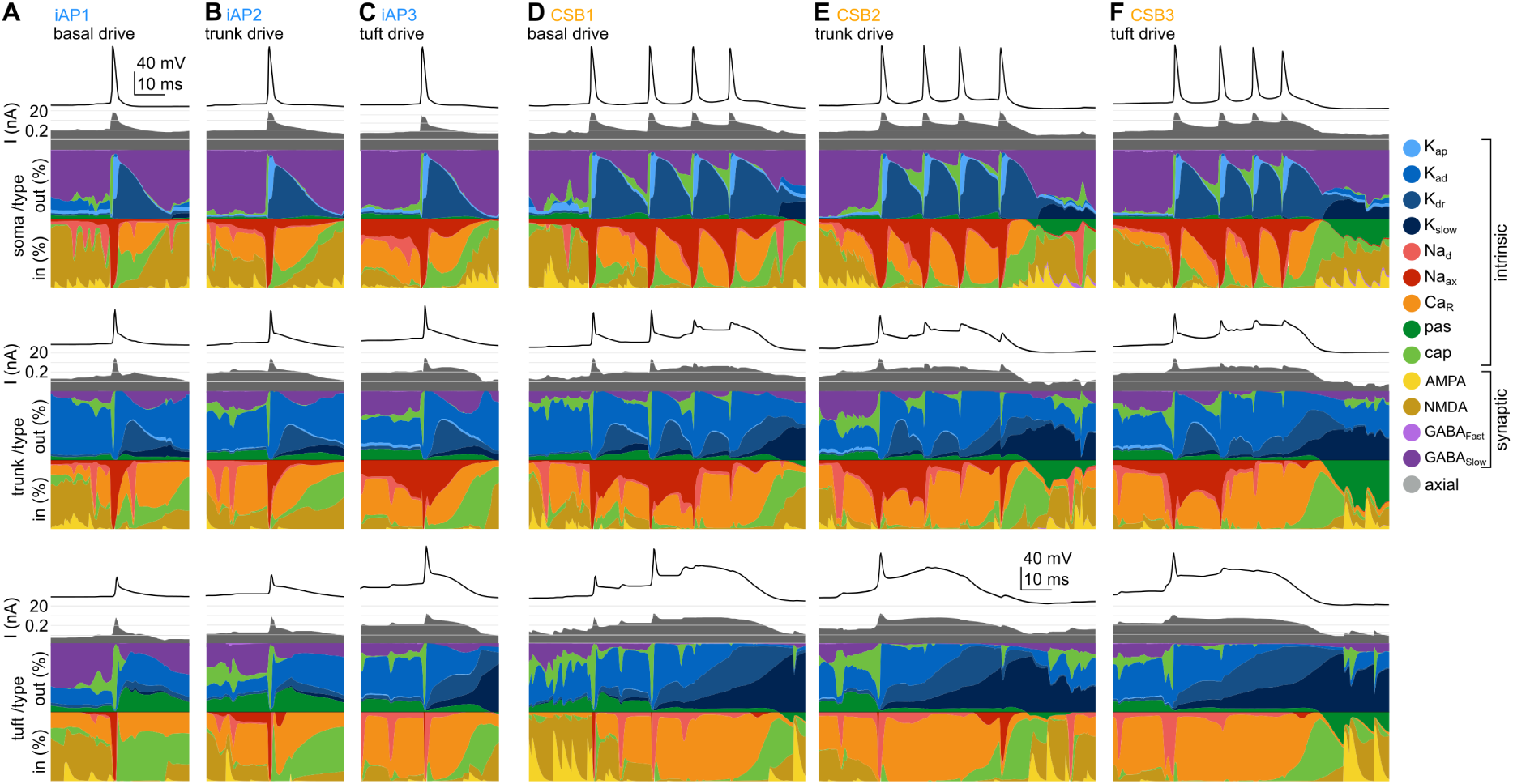
Examples of iAPs and CSBs partitioned by current types. **A-C)** Somatic (top), distal trunk (middle) and tuft (bottom) currents during iAPs partitioned by current type. The same events are shown as in Fig. 8D-F. **D-F**) Somatic (top), distal trunk (middle) and tuft (bottom) currents during CSBs partitioned by current type. The same events are shown as in Fig. 8G-I.

**Fig. S5:**
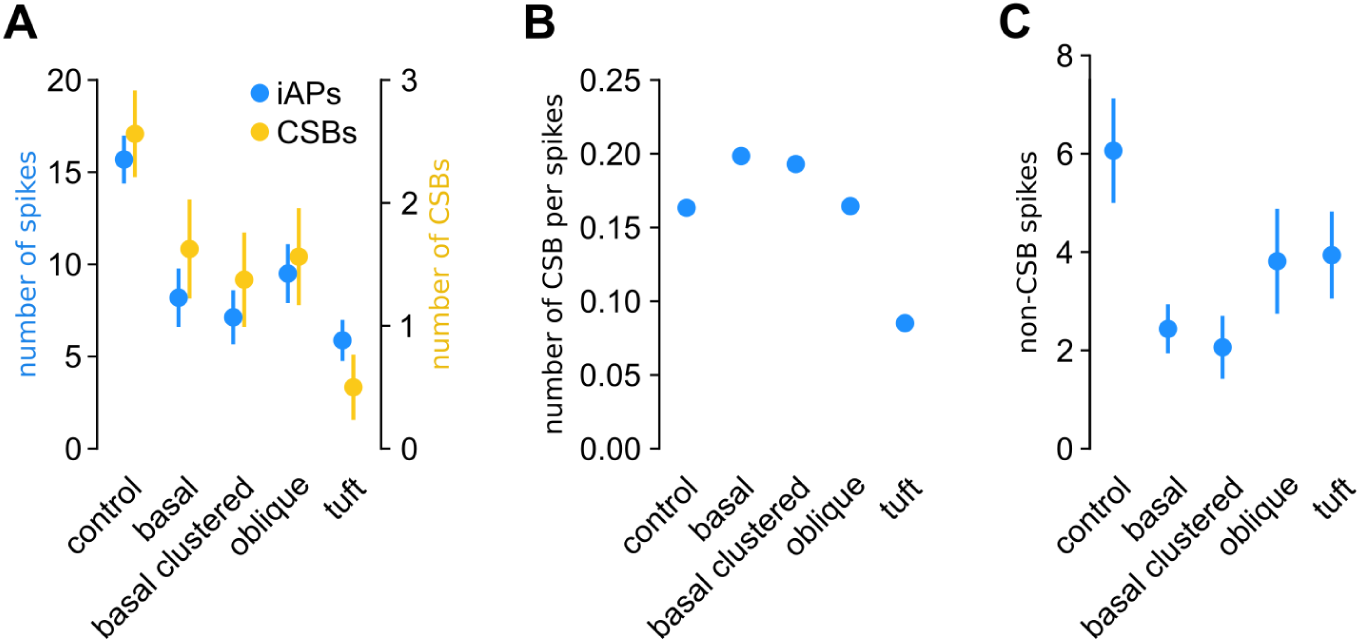
Simulating optogenetic inhibition of inputs targeting different dendritic domains. **A**) The 75% of the input spikes were removed for *N* ≈ 300 randomly selected presynaptic inputs targeting the basal (of the *N*_total_ ≈ 900 inputs), oblique (*N*_total_ ≈ 700) or tuft (*N*_total_ ≈ 400) dendrites. In the *basal clustered* case, the number of synapses in the basal input clusters affected by the inhibition were matched to the number of synapses in tuft input clusters. These manipulations significantly reduced both the average number of output spikes and the complex spike bursts (CSBs) per lap (Wilcoxon signed-rank test compared to control, p < 0.01 in all cases), with the tuft inhibition having the strongest effect on both spikes and CSBs. Moreover, inhibiting the tuft was more specific than that of the oblique of basal domains as it reduced the number of CSBs to a greater extent than the number of spikes. **B**) The number of CSB events per spikes is most strongly reduced by tuft inhibition. **C**) Simulated optogenetic inhibition also reduced the number of spikes outside of CSBs. This effect was strongest for basal dendrites. Symbols show mean across 16 laps, error bars indicate SE.

**Fig. S6:**
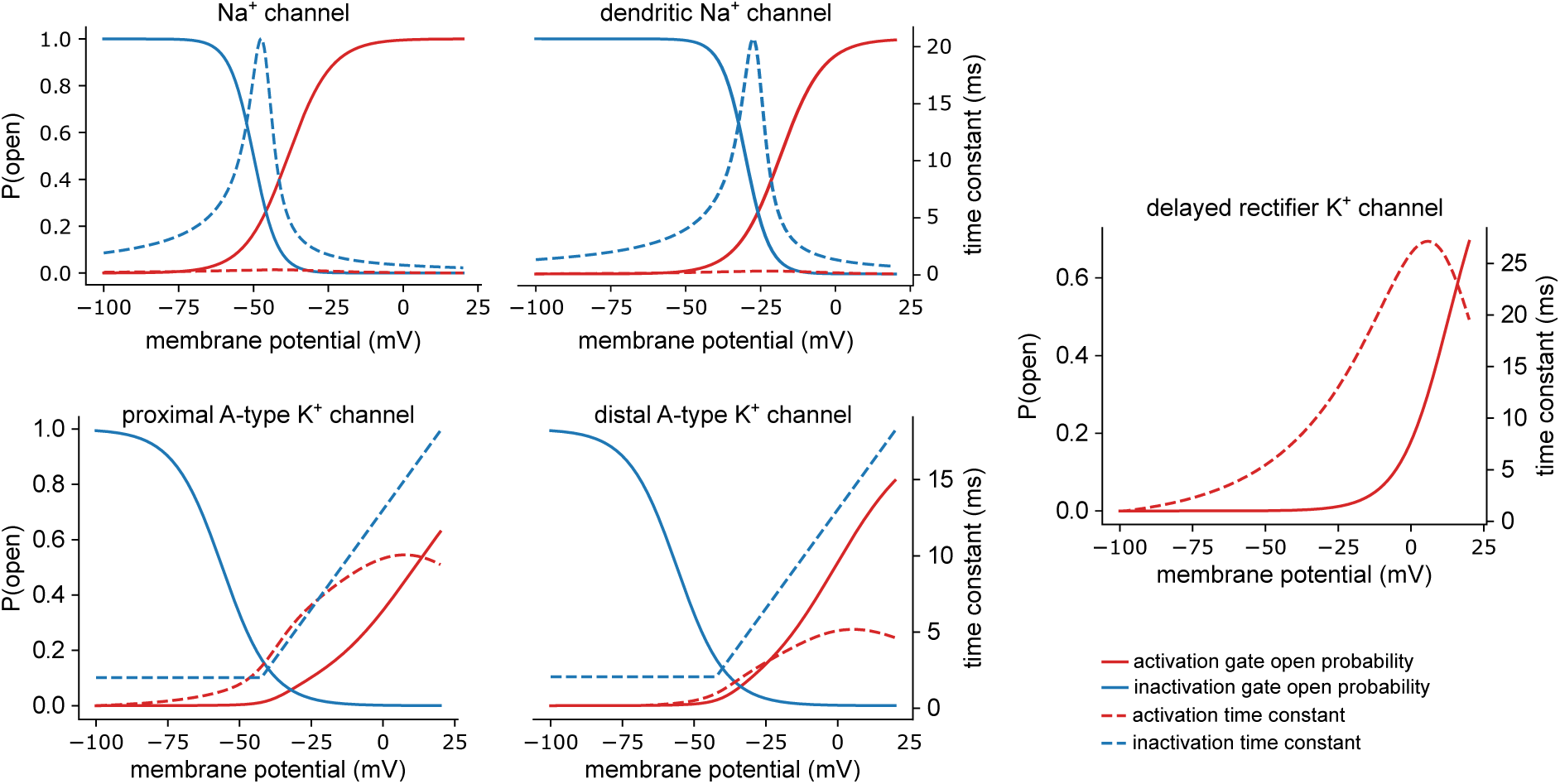
Steady-state activation (solid red line) and inactivation (solid blue line) curves of the voltage-gated ion channels expressed in the detailed model (left axis). Dashed lines indicate the time constants of the channels (right axis).

**Fig. S7:**
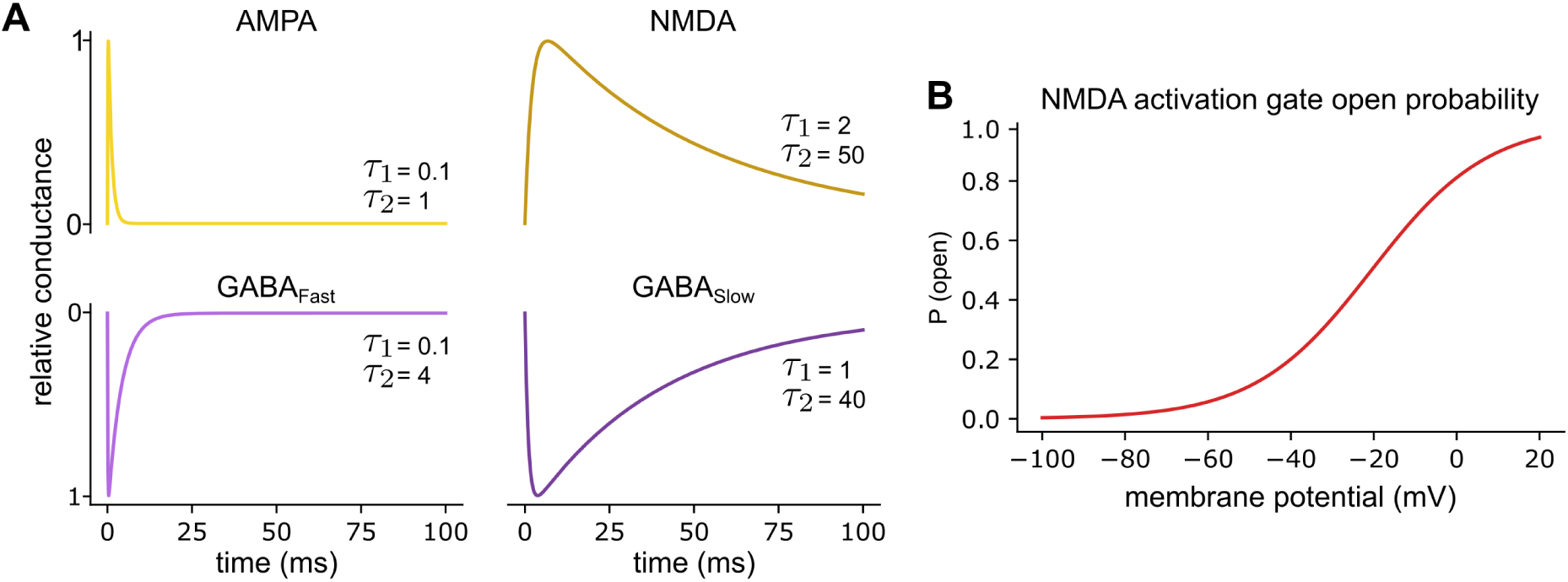
Summary of synaptic channel kinetics. **A**: Time course of excitatory (red) and inhibitory (blue) postsynaptic potentials. We used double exponential kinetics with rise time (*τ*_1_) and decay time (*τ*_2_) constants presented in milliseconds. The reversal potential of the synaptic currents was 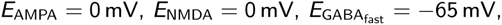 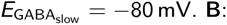 Steady-state activation curve of the NMDA channel.

